# Sec14L6 is a PS and PI4P transporter that promotes lipid droplet formation

**DOI:** 10.1101/2024.10.20.619318

**Authors:** Tiantian Zhou, Yuanjiao Du, Xuewen Hu, Wei-Ke Ji

## Abstract

Lipid droplets (LDs) are evolutionarily conserved organelles that are crucial for cellular metabolism. LD biogenesis and growth occurs in the ER and requires the supply of phospholipid from the ER, but the molecular basis is largely unclear. Here, we have identified Sec14L6, a unique protein of the Sec14 family, as a PS/PI4P transporter required for LD biogenesis. Sec14L6 knockout (KO) greatly reduces the number of LDs that can be rescued by wild-type rather than lipid transfer-defective Sec14L6 mutants. We found that Sec14L6 directly interacts with ACSL3, and that this interaction facilitates targeting of Sec14L6 to LDs and activates the PS transfer activity of Sec14L6 *in vitro*. We also identified PGRMC1, an ER membrane protein, as an adaptor that recruits Sec14L6 to the ER. Furthermore, depletion of Sec14L6 impairs differentiation of adipose-derived mesenchymal stem cells. Together, our study suggests that Sec14L6 transports PS and/or PI4P between the ER and LDs to promote LD formation.

## Introduction

Lipid droplets (LDs) are storage organelles that are at the center of lipid and energy homeostasis^1, 2^. LDs store neutral lipids in a hydrophobic core of triacylglycerol and cholesteryl esters surrounded by a phospholipid monolayer associated with proteins^3, 4^. LDs play an important role in cellular adaptation to energy fluctuations and are also involved in other cellular functions such as protein quality control, regulation of gene transcription, lipid and membrane traffic, and inflammatory responses^5^.

Biogenesis of LDs occurs in the endoplasmic reticulum (ER) in response to an excess of neutral lipids following biosynthesis and deposition of neutral lipids in the hydrophobic region of the bilayer^6^. The neutral lipids condense and form the core of a nascent LD, which grows and then buds off the ER to become a mature LD^7^. During LD biogenesis, LDs acquire surface phospholipids from the ER to meet requirements during membrane expansion and growth as the phospholipid monolayer expands. During the initial phase of their life, the monolayers of nascent LDs are continuously connected to the cytosolic leaflet of the ER membrane^8^, in which several key factors has been identified, including Seipin^9–11^, DGATs^12^, ACSL3^13^ and FIT2^14^. Next, LDs bud toward the cytosol and detach from the ER to form mature LDs that continue to maintain physical contact with the ER through lipid transfer protein (LTP)-dependent mechanisms^15^, such as ORPs^16^ and VPS13^17, 18^. This physical interaction could promote the supply of phospholipids for the growth of mature LDs from the ER^19^. Of note, it has been shown that key enzymes of phospholipid and sterol biosynthesis could translocate to the LD surface for local lipid synthesis during mature LD growth^20^. The molecular basis of LD biogenesis is not completely defined and the question of whether and how lipid transfer occurs between the ER and LDs is largely unknown.

Sec14-like (Sec14L) proteins constitute an atypical subfamily of phosphoinositide transfer proteins (PITP) that are conserved from yeast (Sec14p) to humans (Sec14L1-6)^21, 22^. Sec14L proteins are involved in diverse physiological functions. Sec14L2 has been reported to modulate cholesterol synthesis and hepatitis C virus replication^23, 24^. In zebrafish, zSec14L3 (a homolog of human Sec14L2) regulates Wnt/Ca^2+^ signaling and zebrafish vasculogenesis^25, 26^. In addition, zSec14L3 promotes the conversion of PI4P to PI3P to control ER-mediated endosomal fission^27^. To date, it is unclear whether and how Sec14L proteins are involved in LD dynamics. In this study, we identified Sec14L6, a unique protein of Sec14 family, as a PS/PI4P transporter that was essential for LD formation and differentiation of adipose-derived mesenchymal stem cells (ADSC). Sec14L6 directly interacted with ACSL3, which facilitated the targeting of Sec14L6 to LDs and stimulated the PS transfer activity of Sec14L6 *in vitro*. We further identified PGRMC1 as an adaptor that recruited Sec14L6 to the ER. Taken together, our results suggest that Sec14L6 transports PS and/or PI4P between the ER and LDs to promote LD formation.

## Results

### The formation of LDs is impaired in Sec14L6-depleted cells

To investigate whether Sec14L proteins are involved in LD dynamics, we performed a candidate RNAi screen. In this screen, HeLa cells were cultured in Earle’s Balanced Salt Solution (EBSS) for approximately 12 hours to deplete existing LDs, followed by stimulation of nascent LD formation by a 2-hour treatment with oleic acid (OA) (EBSS-OA). Suppression of Sec14L proteins, including Sec14L1-L5, by small interfering RNAs (siRNAs) had no significant effect on the number or size of LDs compared to control (Fig. 1A, top and middle; Fig. 1B, C & Fig. S1A-E). In contrast, depletion of Sec14L6 by two independent siRNAs greatly reduced the number of LDs (Fig. 1A, bottom; Fig. S1F, G). In addition, Sec14L6 depletion also appeared to cause LD clustering at the cell periphery (Fig. 1A, bottom).

**Fig. 1.**
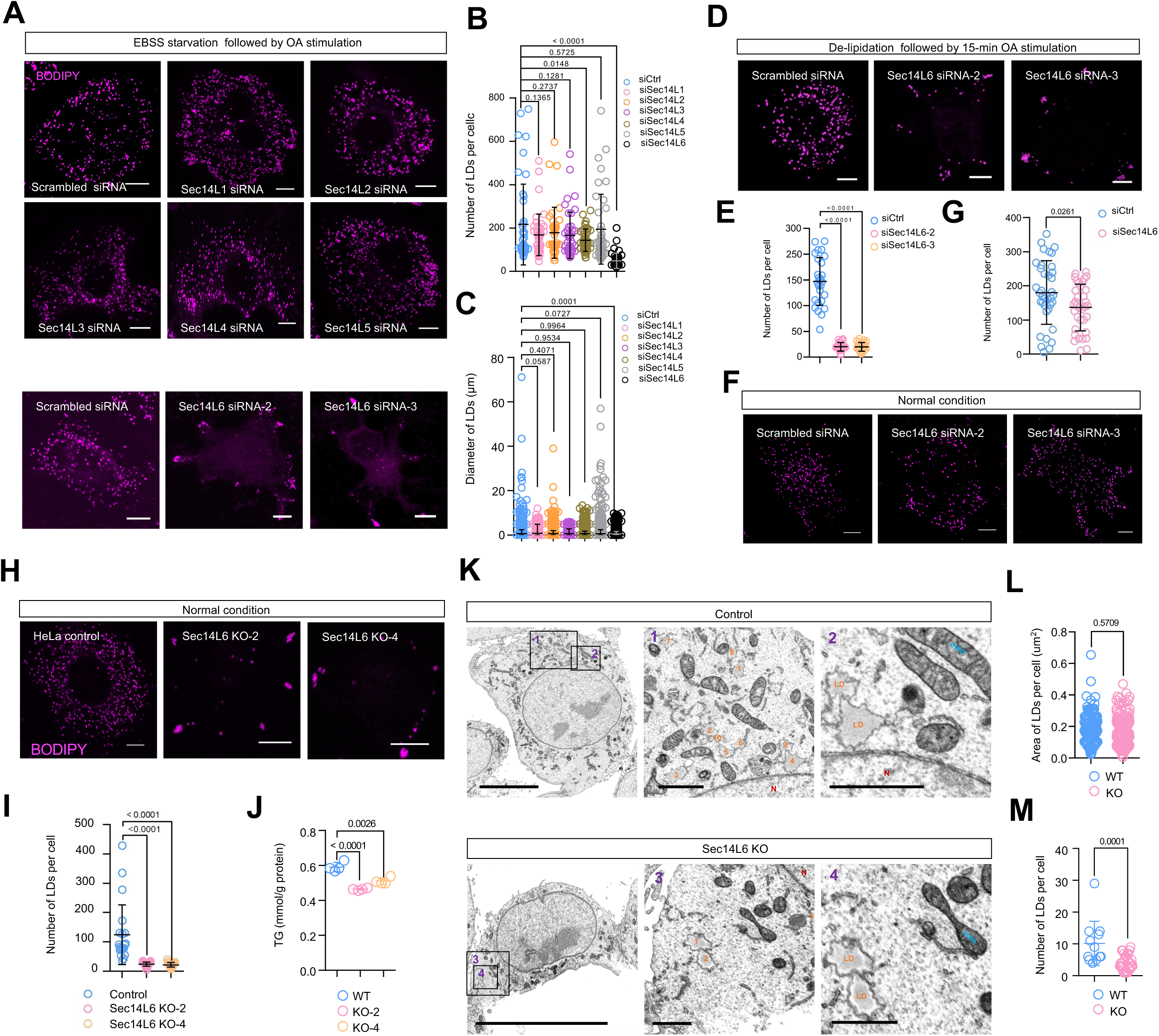
Depletion of Sec14L6 impairs the formation of LDs. **A**. Representative images of live BODIPY (magenta)-labeled HeLa cells treated with scrambled or siRNAs targeting Sec14L1-6 upon EBSS-OA treatments. **B, C**. The number (**B**) or size (**C**) of LDs in cells as shown in (**A**) in more than 3 independent experiments. Ordinary one-way ANOVA with Tukey’s multiple comparisons test. Mean ± SD. **D.** Representative images of live BODIPY (magenta)-labeled HeLa cells treated with scrambled or siRNAs targeting Sec14L6 upon de-lipidation followed by ∼15-min OA treatment. **E.** The number of LDs in cells as shown in (**D**) in 3 independent experiments. Ordinary one-way ANOVA with Tukey’s multiple comparisons test. Mean ± SD. **F.** Representative images of live BODIPY (magenta)-labeled HeLa cells treated with scrambled or siRNAs targeting Sec14L6 under normal culture condition. **G.** LD number in cells as shown in (**F**) in 3 independent experiments. Two-tailed unpaired student t-test. Mean ± SD. **H.** Representative images of live BODIPY (magenta)-labeled control or Sec14L6 KO HeLa cells under normal culture condition. **I.** LD number in cells as shown in (**H**) in more than 3 independent experiments. Ordinary one-way ANOVA with Tukey’s multiple comparisons test. Mean ± SD. **J.** Cellular levels of TG in control or Sec14L6 KO HeLa cells. Measurements of TG were from 3 independent experiments. Ordinary one-way ANOVA with Tukey’s multiple comparisons test. Mean ± SD. **K.** Representative SEM images of control or Sec14L6 KO HeLa cells under normal culture condition. **L, M**. The size (**L**) or number (**M**) or of LDs in cells as shown in (**K**) in 3 independent experiments. Two-tailed unpaired student t-test. Mean ± SD. Scale bar, 10μm in the whole cell images and 2μm in the insets in (A, D, F, H & K).

Treatment with EBSS not only removes lipids from cells but also other nutrients. To confirm a specific role of Sec14L6 in LD formation, we investigated the role of Sec14L6 under canonical LD induction conditions, in which cells were cultured in lipoprotein-free medium followed by OA loading for ∼15 minutes as previously reported^10^. Importantly, Sec14L6 depletion greatly reduced LD number under this condition (Fig. 1D, E). Noteworthy, Sec14L6 depletion did not dramatically affect the number of existing LDs under normal growth condition (Fig. 1F, G), in which LD biogenesis was not strongly stimulated, suggesting a specific role of Sec14L6 in LD formation.

We further confirmed the role of Sec14L6 in LD formation in CRISPR-Cas9-mediated Sec14L6 knockout HeLa cells (Sec14L6-KO; Fig. S1H-K). Consistently, LD formation was severely impaired in two independent Sec14L6-KO clones (KO-2 & −4), as evidenced by a great reduction in LD abundance (Fig. 1H, I). In addition, we observed the clustering of LDs in the periphery of Sec14L6-KO cells, similar to cells treated with Sec14L6 siRNAs. The mechanism through which Sec14L6 affects LD positioning is unclear and requires in-depth investigations in the future.

Notably, siRNA-mediated depletion of Sec14L6 resulted in only a slight decrease in cellular triglycerides (TG) levels, to a similar extent as depletion of known essential factors of LD biogenesis such as Seipin or ACSL3 (Fig. S1L). Accordingly, TG levels were also only slightly reduced in Sec14L6 KO cells (Fig. 1J), suggesting that Sec14L6 regulates LD formation likely independently of TG synthesis.

The clustering of LDs makes it difficult to measure LD size by confocal microscopy, so we performed scanning electron microscopy (SEM). SEM micrographs showed that the size of LDs in Sec14L6 KO cells was not significantly altered (Fig 1K, L). Importantly, SEM confirmed that Sec14L6 KO resulted in a significant reduction in LD abundance compared to control (Fig 1K, M).

Next, we investigated the functional relationship between Seipin and Orp5^28, 29^, key factors in LD biogenesis, and Sec14L6. Overexpression of GFP-Seipin (Fig. S2A) or GFP-Orp5 (Fig. S2B) failed to restore the LD defect in Sec14L6 KO line (Fig. S2C, D). Meanwhile, GFP-Sec14L6 did not rescue the LD phenotype in Orp5- (Fig. S2E, F, H, K-M) or Seipin- (Fig. S2G, I, K, L, N) depleted HeLa cells. In addition, overexpression of Sec14L6 did not strongly alter LD size and number (Fig. S2J, K & L). These results suggest a specific role of Sec14L6 in LD biogenesis. Collectively, our results suggest that Sec14L6, but not other Sec14L proteins, is required for LD formation.

### ACSL3 facilitates the association of Sec14L6 with LDs

We next investigated whether Sec14L6 associates with LDs. Sec14L6 tagged with GFP at its N-terminus (GFP-Sec14L6, Fig. 2A) or C-terminus (Sec14L6-GFP, Fig. S3A) was soluble in the cytosol without membrane associations under OA treatment.

**Fig. 2.**
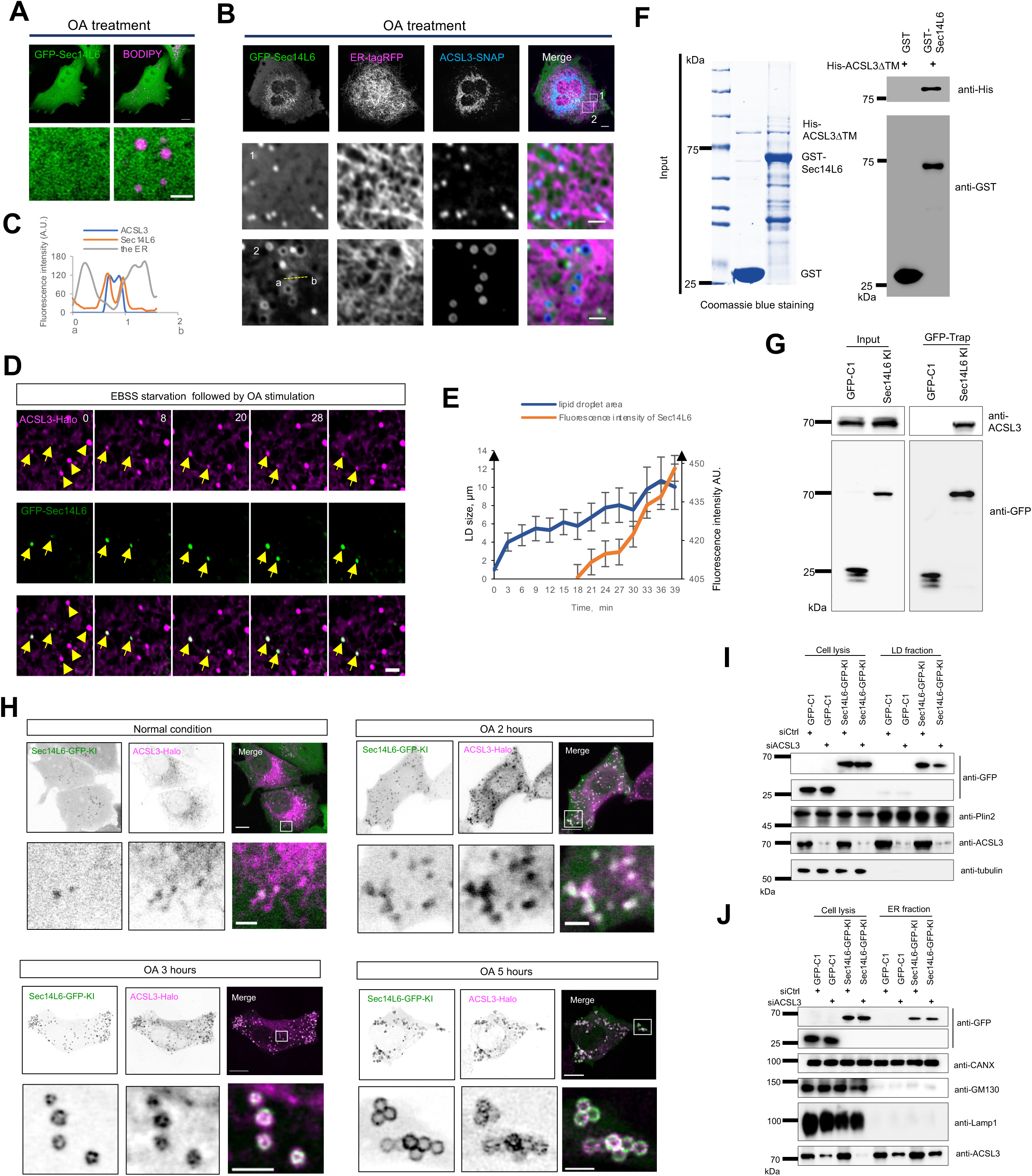
ACSL3 facilitates the association of Sec14L6 with LDs. **A.** Representative images of a live BODIPY (magenta)-labeled HeLa cell transiently transfected with GFP-Sec14L6 (green) under OA stimulation with an inset on the bottom. **B.** Representative images of a live HeLa cell transiently transfected with GFP-Sec14L6 (green), ER-tagRFP (magenta) and ACSL3-SNAP (blue) under OA stimulation (>6 h) with two insets on the bottom. A yellow arrow denotes a Sec14L6-decorated LD. **C.** Line-scan analysis of the bottom inset from (**B**). **D.** Time-lapse montages of a HeLa cell transiently transfected with GFP-Sec14L6 (green) and ACSL3-mCh (magenta) with arrows denoting growing LDs. **E.** The correlation between LD diameter and GFP-Sec14L6 fluorescence intensity as in (**J**). **F.** GST pulldown assays showed that GST-Sec14L6 was pelleted with purified His-ACSL3-ΔTM *in vitro*. The Coomassie blue staining of proteins used in the assay was shown on the left. **G.** GFP-Trap assays demonstrated interactions between endogenous Sec14L6-GFP-KI and endogenous ACSL3. **H.** Representative images of live BODIPY-stained Sec14L6-GFP-KI (green) cells transiently transfected with ACSL3-Halo (magenta) in absence or presence of OA loading for indicated time (2, 3 and 5 hours) with insets on the bottom. **I.** Membrane fractionation showing the level of Sec14L6-GFP-KI was reduced in LD fractions upon ACSL3 depletion under LD-induction condition as in Fig. 1D. **J.** Immunoblots showed the level of Sec14L6-GFP-KI in ER membrane fractions of either control or ACSL3-depleted cells using GFP vector as a negative control under LD-induction condition as in Fig. 1D. Scale bar, 10μm in the whole cell images and 2μm in the insets in (A, B, D & H).

Interestingly, we found that ACSL3, the important factor in LD biogenesis^13^, facilitated the recruitment of GFP-Sec14L6 to the surfaces of both small (middle panel) and large LDs (bottom panel) after OA stimulation (Fig. 2B). These small LDs marked GFP-Sec14L6 appeared to be adjacent to the ER. In contrast, line-scan analyses showed that large LDs decorated with GFP-Sec14L6 were not closely associated with the ER (Fig. 2C). These results suggest that ACSL3 specifically facilitates the targeting of Sec14L6 to LDs.

Next, we tracked the process of nascent LD growth in live HeLa cells using time-lapse imaging. GFP-Sec14L6 was colocalized with small LDs marked by ACSL3-Halo, and these small LDs grew as Sec14L6 puncta became bigger (Fig. 2D, E; Video 1).

We then investigated whether Sec14L6 interacts with ACSL3. Indeed, co-IP assays showed that GFP-Sec14L6 interacts with endogenous ACSL3 (Fig. S3B).

To investigate whether or not the Sec14L6-ACSL3 interaction is direct, we performed GST pulldown assays. His-tagged ACSL3 lacking of its transmembrane domain (ACSL3-ΔTM) bound to GST-tagged Sec14L6 but not to the GST tag (Fig. 2F), indicating that Sec14L6 binds to ACSL3.

To avoid potential artifacts of overexpression, we labeled endogenous Sec14L6 with monomeric superfolder GFP (sfGFP) at its C-terminus using Crispr-Cas9 in HeLa cells (Sec14L6-GFP-KI) (Fig. S3C-E). We chose to tag Sec14L6 at the C-terminus in the KI line for two reasons. First, the best sgRNA for KI was located in this region (Fig. S3C). Second, tagging at the C-terminus did not affect the Sec14L6 localization, its interactions with other proteins identified in this study (Fig. S3F, G), or its function in LD formation (Fig. 8D, E).

The Sec14L6-ACSL3 interaction was confirmed at the endogenous level by coIP assays in Sec14L6-GFP KI HeLa cells (Fig. 2G).

Importantly, Sec14L6-GFP-KI was recruited to LDs labeled with ACSL3-mCh even without OA treatment, and the extent of recruitment to LD surface was increased as LDs growed induced by OA stimulation for indicated time (Fig. 2H).

To further investigate whether ACSL3 is required for LD localization of Sec14L6, we performed membrane fractionation. siRNA-mediated depletion of ACSL3 reduced the amounts of endogenous Sec14L6-GFP-KI in LD fractions (Fig. 2I). In contrast, ACSL3 depletion did not substantially alter the level of Sec14L6-GFP-KI in ER fractions (Fig. 2J). Taken together, our results indicated that ACSL3 binds to Sec14L6 and promotes the association of Sec14L6 with LDs.

### Molecular mechanism of the Sec14L6-ACSL3 interaction

Next, we investigated the mechanism underlying the interaction between Sec14L6 and ACSL3. Sec14L6 contains an N-terminal region (NT), a Carl-Trio domain, and a Golgi dynamics domain in the C-terminal region (CT) (Fig. 3A)^21^. Pull-down assays demonstrated that the Carl-Trio domain and the CT region, but not the NT region, could bind to purified His-ACSL3-ΔTM. Of note, the Carl-Trio bound to His-ACSL3-ΔTM to a much higher extent compared to the CT region (Fig. 3B).

**Fig. 3.**
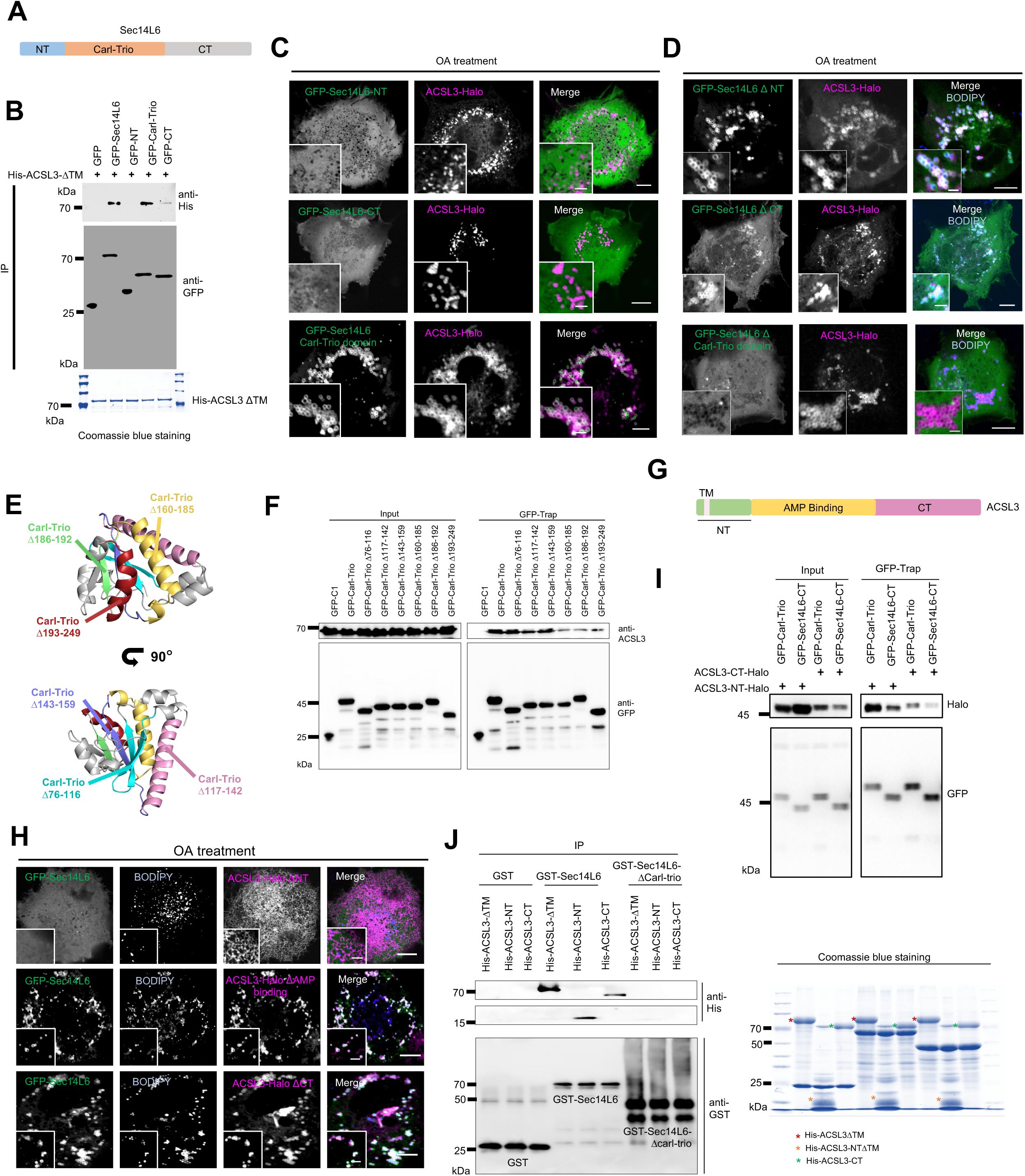
Molecular mechanism underlying the Sec14L6-ACSL3 interaction. **A.** Domain organization of Sec14L6. **B.** Pulldown assays demonstrated that the Carl-Trio domain and CT of Sec14L6 was co-pelleted with purified His-ACSL3ΔTM *in vitro*. The Coomassie blue staining of proteins used in the assay was shown on the bottom. **C.** Representative images of live BODIPY (magenta)-labeled HeLa cells transiently transfected with GFP-Sec14L6 truncations (green; the NT, the Carl-Trio and the CT region) and ACSL3-Halo (magenta) with insets under OA stimulation. **D.** Representative images of live BODIPY (blue)-stained HeLa cells transiently transfected with GFP-Sec14L6 deletion mutants (green; ΔNT, ΔCarl-Trio and ΔCT region) and ACSL3-Halo (magenta) with insets under OA stimulation. **E.** AlphaFold-predicted structure of the Carl-Trio domain of Sec14L6 with secondary structures labeled in different color. **F.** GFP-Trap assays demonstrated interactions between GFP-Sec14L6 truncation mutabnts lacking of these secondary structures as in (**E**) and endogenous ACSL3. **G.** Domain organization of ACSL3. **H.** Representative images of live BODIPY (blue)-stained HeLa cells transiently transfected with GFP-Sec14L6 (green) and ACSL3-Halo deletion mutants (magenta; ΔNT, ΔAMP binding and ΔCT region) with insets under OA stimulation. **I.** GFP-Trap assays demonstrated ACSL3-NT preferentially interacted with Sec14L6-Carl-Trio domain. **J.** GST pulldown assays demonstrated that the Carl-Trio domain of Sec14L6 was co-pelleted with purified His-ACSL3-ΔTM, His-ACSL3-NT and His-ACSL3-CT *in vitro*. The Coomassie blue staining of proteins used in the assay was shown on the right. Scale bar, 10μm in the whole cell images and 2μm in the insets in (C, D & H).

Consistently, live-cell imaging showed that the Carl-Trio domain, but not the NT or CT, colocalized with ACSL3 (Fig. 3C). In addition, deletion of the Carl-Trio domain, but not the NT or CT region, abolished the colocalization between Sec14L6 and ACSL3 (Fig. 3D), suggesting that the Carl-Trio domain is responsible for binding to ACSL3 on LD surface.

We then dissected the Carl-Trio domain to identify the region responsible for the interaction with ACSL3. We found that deletion of a helix (residues 160-185), a sheet (residues 186-192) or a helix (residues 193-249) partially abolished the interaction with endogenous ACSL3, whereas deletion of other secondary structures in this domain had no effect (Fig. 3E, F), suggesting that the region of the Carl-Trio domain containing the helix-sheet-helix is required for the interaction with ACSL3.

ACSL3 has a TM domain in the NT region, an AMP-binding domain and a CT region (Fig. 3G). Deletion of the NT region of ACSL3 (the TM domain is retained), but not the AMP-binding domain or the CT, abolished colocalization between ACSL3 and Sec14L6 (Fig. 3H), suggesting that the NT region of ACSL3 is required to facilitate Sec14L6 recruitment to LDs.

In addition, GFP-Trap assays revealed multivalent interactions between the domains of Sec14L6 and ACSL3, with the interaction between Sec14L6-Carl-Trio and ACSL3-NT being the most prominent (Fig. 3I). Accordingly, GST pulldown assays demonstrated that GST-Sec14L6 could bind to His-ACSL3-NT-ΔTM and His-ACSL3-CT, albeit to a much lesser extent than ACSL3-ΔTM. Notably, GST-Sec14L6-Δcarl-trio did not bind to purified ACSL3-ΔTM or its truncations (Fig. 3J).

Interestingly, we found that Sec14L6 could self-interact (Fig. S4A), and the Carl-Trio domain and the CT region of Sec14L6 contributed to the self-interaction (Fig. S4B, C). Cross-linking mediated by disuccinimidyl suberate (DSS) promoted oligomerization of Flag-Sec14L6 in HeLa cells in a dose-dependent manner (Fig. S4D). Collectively, our results suggested that the Sec14L6-ACSL3 interaction is mediated by multivalent interactions between Sec14L6 oligomers and ACSL3 and that the recruitment of Sec14L6 to LDs depends on the interaction between the Carl-Trio domain of Sec14L6 and the NT of ACSL3.

### The Carl-Trio domain of Sec14L6 targets LD via an AH

Surprisingly, we found that, unlike the full-length Sec14L6, the Carl-Trio domain was able to target LDs in the absence of ACSL3 (Fig. 4A). Remarkably, although ACSL3 was able to facilitate the association of full-length Sec14L6 with LDs, depletion of ACSL3 did not block the LD localization of the Carl-Trio domain after OA loading (Fig. 4B, C).

**Fig. 4.**
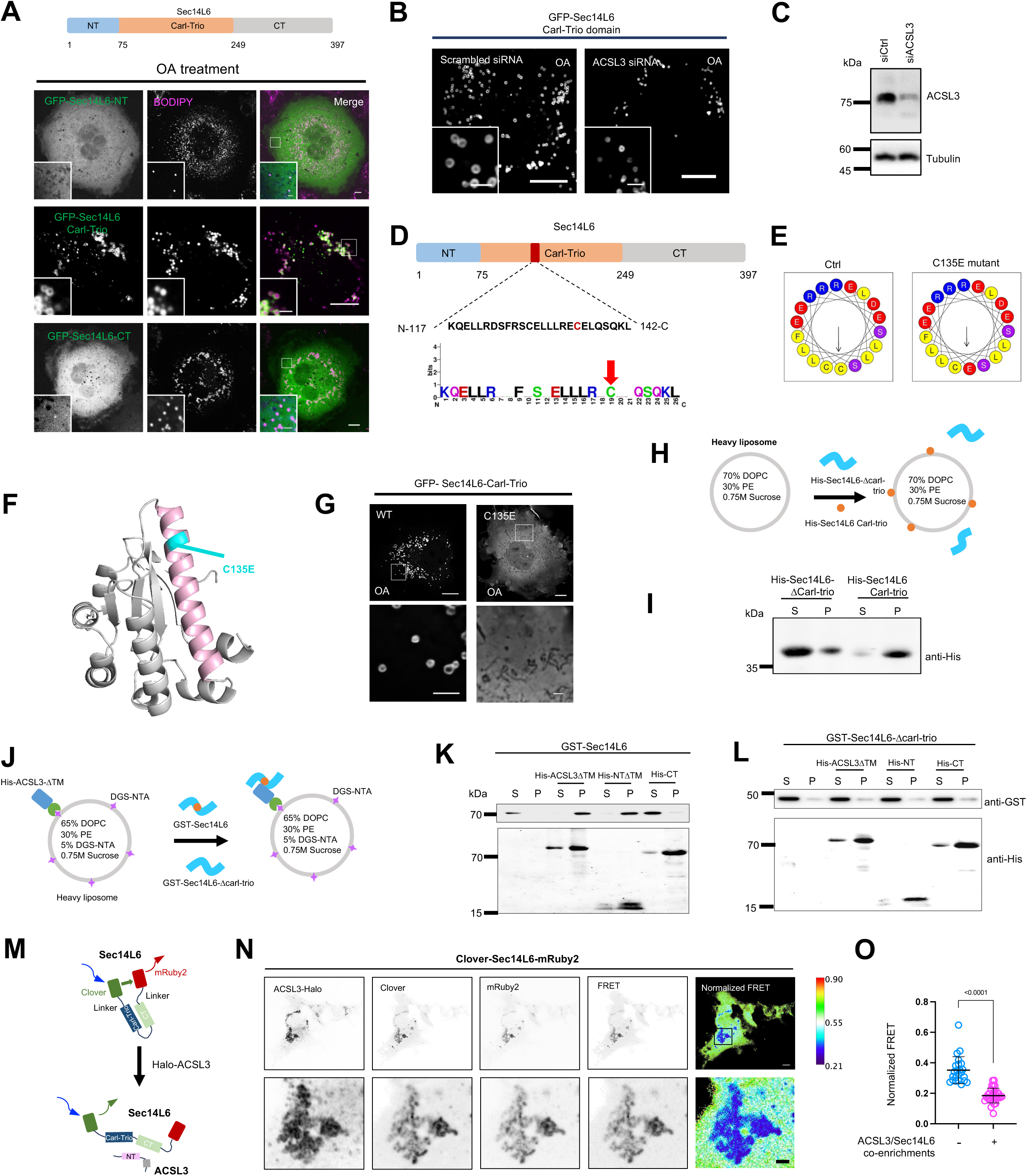
The Carl-Trio domain of Sec14L6 targets LD via an AH. **A.** Representative images of live BODIPY (magenta)-stained HeLa cells transiently expressing GFP-Sec14L6 truncations (green; the NT, the Carl-Trio and the CT region) under OA stimulation. **B.** Representative images of live HeLa cells transiently expressing GFP-Sec14L6-Carl-Trio upon scrambled or ACSL3 siRNAs treatments under OA stimulation. **C.** Immunoblots showing the efficiency of ACSL3 siRNAs used in (**F**). **D, E**. Schematic representation of an AH in the Carl-Trio domain of Sec14L6 with a point mutation C135E (**D**) that would impair the AH (**E**). **F**. AlphaFold-predicted structure of the Carl-Trio domain of Sec14L6 showing the C135E (cyan) in the AH (magenta). **G.** Representative images of live HeLa cells transiently expressing GFP-Sec14L6-Carl-Trio and GFP-Sec14L6-Carl-Trio-C135E under OA stimulation. **H, I.** Liposome pelleting assays showing His-Sec14L6-Carl-trio (0.25 μM), but not His-Sec14L6-ΔCarl-Trio (0.25 μM), was co-pelleted with sucrose-loaded liposomes. S, supernatant; P, pellet. **J, K.** Liposome pelleting assays showing the addition of His-ACSL3ΔTM (0.25 μM) and His-ACSL3-NTΔTM (0.25 μM), but not His-ACSL3-CT (0.25 μM), promotes the membrane association of GST-Sec14L6 (0.25 μM). **L.** As in (**K**), the addition of His-ACSL3ΔTM (0.25 μM) or the two ACSL3 truncations (0.25 μM for each mutant) had no effect on the membrane association of GST-Sec14L6ΔCarl-trio. **M.** Schematic diagram of Clover-Sec14L6-mRuby2 protein. **N.** Confocal images showing live HeLa cells transiently expressing Clover-Sec14L6-mRuby2 along with ACSL3-Halo with insets on the bottom. Normalzed FRET signal is indicated by a pseudocolored image. **O.** Normalzed FRET signal from either ACSL3/Sec14L6 co-enriched regions or other cells without substabtial co-enrichments in HeLa cells in 3 independent experiments. Each dot represents one normalized FRET signal ratio from one region. Two-tailed unpaired student t-test. Mean ± SD. Scale bar, 10μm in the whole cell images and 2μm in the insets in (A, B, G & N).

The Carl-Trio domain of MOSPD2 has been shown to target LD via an amphipathic helix (AH)^30^. We thus hypothesized that the Carl-Trio domain of Sec14L6 might target LDs via the same mechanism. A segment in the Carl-Trio domain of Sec14L6 was predicted to fold as an AH (Fig. 4D, E). Mutation of a conserved residue (C135) on the hydrophobic side of this AH reduced the localization on LDs (Fig. 4F, G), indicating that the AH is required for the localization of Sec14L6-Carl-Trio to the surface of LDs.

To further validate these results, we performed *in vitro* liposome pelleting assays (Fig. 4H). The his-tagged Carl-Trio domain of Sec14L6, but not His-Sec14L6-ΔCarl-Trio, could be substantially pelleted by sucrose-loaded liposomes (Fig. 4I), indicating that GST-Carl-Trio can target liposome membranes *in vitro*. In contrast, purified GST-Sec14L6 was barely associated with liposomes (Fig. 4J, K). Importantly, the addition of purified His-ACSL3-ΔTM significantly enhanced the association of Sec14L6 with liposomes. The effect was mediated by ACSL3-NT, as the addition of His-ACSL3-NT-ΔTM, but not His-ACSL3-CT, promoted the targeting of GST-Sec14L6 to liposomes (Fig. 4K). Importantly, neither His-ACSL3-ΔTM nor the two truncations had any effect on liposome association of GST-Sec14L6-ΔCarl-trio (Fig. 4L). Therefore, our results indicated that the Carl-Trio domain of Sec14L6, but not the full-length Sec14L6 protein, is able to target membranes.

Since the AH was not located in the region required for interaction with ACSL3 (Fig. 3E, F), we therefore speculate that ACSL3 may trigger conformational changes of Sec14L6 to release the self-inhibitory configuration. To test this idea, we performed fluorescence resonance energy transfer (FRET) assays to monitor Sec14L6 conformational changes as previously reported^31^. We fused the fluorescent proteins Clover and mRuby2 to the N and C termini of Sec14L6, respectively, and collected the fluorescent signals from Clover-Sec14L6-mRuby2 in HeLa cells (Fig. 4M). Importantly, at LD regions with extensive recruitment of Clover-Sec14L6-mRuby2 by Halo-ACSL3, normalized FRET signals decreased significantly than other region without substantial co-enrichments (Fig. 4N, O). These results indicated that ACSL3 overexpression caused an increase in the distance between Clover and mRuby2, highlighting the role of ACSL3 in the release of the Sec14L6 self-inhibitory conformation.

### PGRMC1 recruits Sec14L6 to the ER

As a potential lipid transporter involved in LD formation, we hypothesized that Sec14L6 may be associated with the ER. To investigate whether and how Sec14L6 recognizes the ER, we performed mass spectrometry (MS) to identify Sec14L6 interacting proteins on the ER. To this end, we identified PGRMC1 (Fig. 5A). PGRMC1 is a component of a progesterone-binding protein complex^32^ and has been reported to play a role in heme homeostasis^33^ and cholesterol synthesis^34^. coIP assays confirmed that Sec14L6 interacted with PGRMC1 at endogenous levels in Se14L6-GFP-KI line (Fig. 5B).

**Fig. 5.**
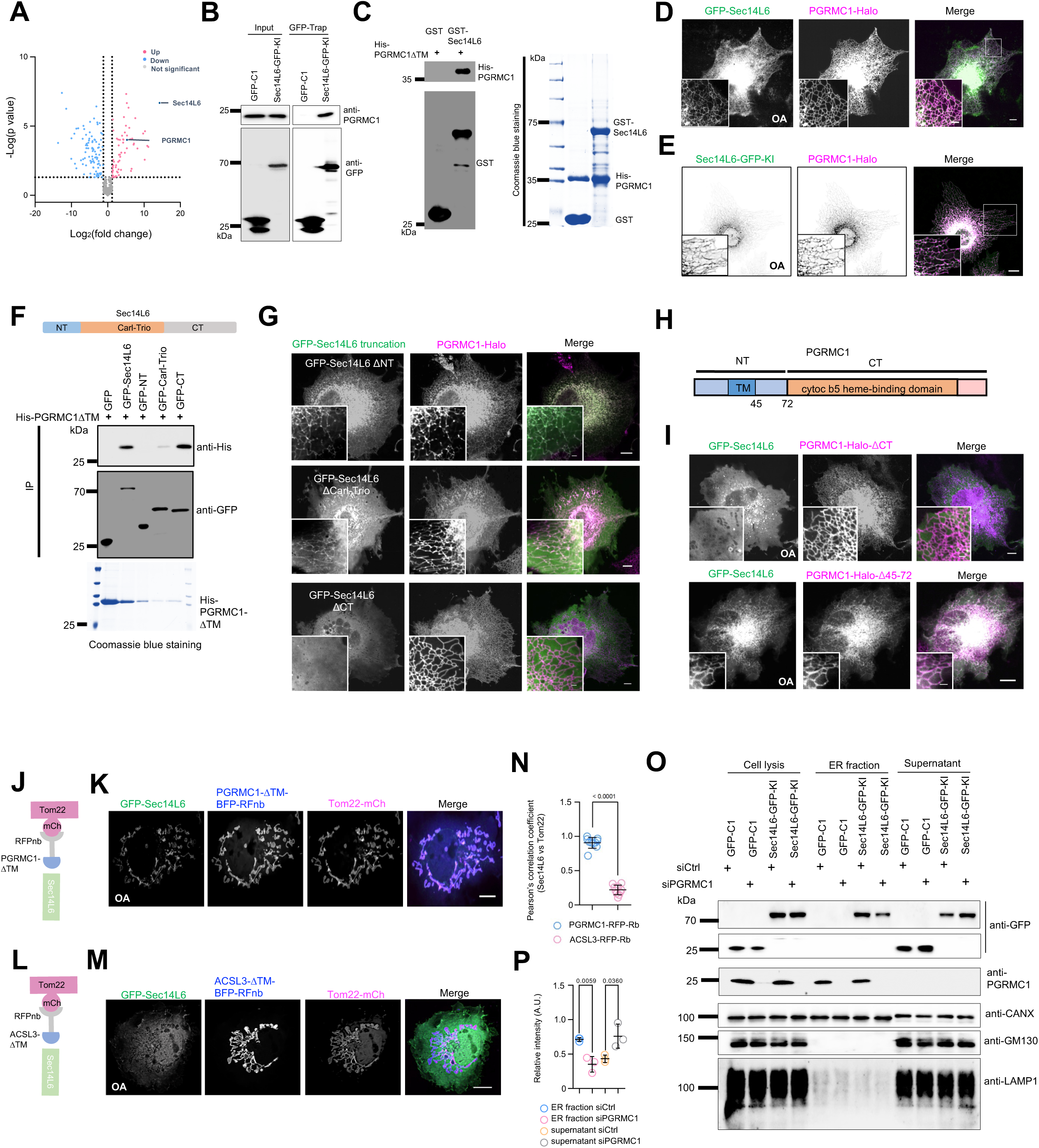
PGRMC1 recruits SEC14L6 to the ER. **A.** Volcano plot of protein candidates coIPed with GFP-Sec14L6 in HEK293 cells compared with protein candidates coIPed with GFP tag only. After removal of proteins that coIPed with GFP tag, candidates that were considered significant (−log [P value] > 1.3; P< 0.05) were labeled in orange (Log2 [fold change] > 0; increased in abundance) or blue (Log2 [fold change] < 0; decreased in abundance). **B.** GFP-Trap assays demonstrated interactions between endogenous Sec14L6-GFP-KI and endogenous PGRMC1. **C.** GST pulldown assays demonstrated that purified GST-Sec14L6, but not GST tag, was pelleted with purified His-PGRMC1 in vitro. The Coomassie blue staining of proteins used in the assay was shown on the right. **D.** Representative images of a live HeLa cells transiently transfected with GFP-Sec14L6 (green) and PGRMC1-Halo (magenta) with one boxed region on the bottom upon OA treatments. **E.** Representative images of live BODIPY-stained Sec14L6-GFP-KI (green) cells transiently transfected with PGRMC1-Halo (magenta) in presence of OA loading with insets. **F.** Pulldown assays demonstrated that the CT of Sec14L6 was pelleted with purified His-PGRMC1 in vitro. The Coomassie blue staining of proteins used in the assay was shown on the bottom. **G.** Representative images of live HeLa cells transiently transfected with indicated GFP-Sec14L6 truncations (green) and PGRMC1-Halo (magenta) with insets upon OA treatments. **H.** Domain organization of PGRMC1. **I.** Representative images of live HeLa cells transiently transfected with GFP-Sec14L6 (green) and indicated PGRMC1-Halo truncations (magenta) with insets upon OA treatments. **J.** Schematic cartoon of RFPnb-mediated recruitment of PGRMC1-BFP-RFnb to the outer mitochondrial membranes (Tom22-mCh) in presence of GFP-Sec14L6. **K.** Representative images of a HeLa cell expressing PGRMC1-BFP-RFnb (blue), GFP-Sec14L6 (green) and Tom22-mCh (magenta). **L.** Schematic cartoon of RFPnb-mediated recruitment of ACSL3-BFP-RFnb to the outer mitochondrial membranes (Tom22-mCh) in presence of GFP-Sec14L6. **M.** Representative images of a HeLa cell expressing ACSL3-BFP-RFnb (blue), GFP-Sec14L6 (green) and Tom22-mCh (magenta) **N.** Pearson’s correlation coefficient of GFP-Sec14L6 colocalized with Tom22-mCh either in presence of PGRMC1-BFP-RFnb (22 cells) or ACSL3-BFP-RFnb (25 cells) of BFP-RFPnb-RHOBTB3. Each dot represents a value of pearson’s correlation coefficient of a cell. More than 3 independent experiments. Two-tailed unpaired student t-test. Mean ± SD. **O.** Immunoblots showed the level of Sec14L6-GFP-KI in ER membrane fractions of either control or PGRMC1-depleted cells stably using GFP vector as a negative control. **P.** Ratio of Sec14L6-GFP-KI level of ER fractions between control and PGRMC1 depletion based on 3 independent assays. Two-tailed unpaired student t-test. Mean ± SD. Scale bar, 10μm in the whole cell images and 2μm in the insets in (D, E, G, I, K & M).

Next, we performed *in vitro* pull-down assays to investigate whether Sec14L6 binds to PGRMC1. In this assay, purified GST-Sec14L6, but not the GST tag alone, bound to His-PGRMC1-ΔTM (Fig. 5C).

Importantly, live-cell imaging showed that cytosolic GFP-Sec14L6 was recruited to the ER marked by PGRMC1-Halo (Fig. S4E), and the treatment with OA appeared to promote the recruitment (Fig. 5D). Interestingly, coIP assays showed that the Sec14L6-PGRMC1 interaction was not strongly affected by OA (Fig. S4F), suggesting that OA may promote the recruitment in an indirect manner.

In addition, Sec14L6 could not be recruited to the ER by PGRMC2, a PGRMC1 homolog that is critical for adipocyte function^35^ (Fig. S4G), suggesting a specific interaction between Sec14L6 and PGRMC1. Importantly, live-cell imaging confirmed that endogenous Sec14L6-GFP-KI could be recruited to the ER by PGRMC1-Halo (Fig. 5E).

We then asked how Sec14L6 interacts with PGRMC1. Pull-down assays showed that the CT region of Sec14L6 bound to His-PGRMC-ΔTM to a much higher extent compared to the Carl-Trio or the NT region (Fig. 5F). Accordingly, live-cell imaging showed that deletion of the CT, but not the NT or the Carl-Trio domain, abolished the association with PGRMC1 (Fig. 5G), indicating that the CT is required for interaction with PGRMC1.

PGRMC1 possessed a TM domain in its NT and a cytochrome b5 heme-binding domain in the CT region (Fig. 5H). Deletion of the CT of PGRMC1, but not the NT (amino acids 45-72; the TM is retained), abolished the colocalization between PGRMC1 and Sec14L6 (Fig. 5I), indicating that the CT of PGRMC1 is required for Sec14L6 recruitment. Collectively, our results suggested that the recruitment of Sec14L6 to the ER is mediated by Sec14L6-CT and PGRMC1-CT.

To further validate this recruitment, we performed knock-sideway assays. Utilizing a specific interaction between RFP and RFP nanobody (RFPnb), we ectopically targeted PGRMC1-ΔTM-RFPnb to the outer mitochondrial membranes (OMM) via Tom22-mCh (Fig. 5J). Remarkably, cytosolic GFP-Sec14L6 was strongly recruited to the OMM that were positive for PGRMC1 (Fig. 5K, N). In contrast, GFP-Sec14L6 was unable to be recruited to the OMM by ACSL3 (Fig. 5L-N), suggesting that ACSL3 plays a role in regulating Sec14L6 configuration other than a recruitment factor. These results suggest that PGRMC1 functions as an adaptor to recruit Sec14L6 to the ER.

We then asked whether PGRMC1 is required for targeting Sec14L6 to the ER. Subcellular fractionation showed that suppression of PGRMC1 by siRNAs significantly reduced the level of Sec14L6 in the ER fractions of Sec14L6-GFP-KI (Fig. 5O, P) or HeLa cells transiently expressing GFP-Sec14L6 (Fig. S4H). In addition, PGRMC1 was not required for the LD targeting of Sec14L6, as live-cell images showed that PGRMC1 depletion did not abolish the LD localization of Sec14L6 in the Sec14L6-GFP-KI line (Fig. S4I). Overall, our results demonstrated that PGRMC1 recruits Sec14L6 to the ER.

### Sec14L6 transfers PS and PI4P *in vitro*

Next, we asked how Sec14L6 promotes LD biogenesis. Sec14L6 belongs to the Sec14 protein family of PITPs, which have been shown to transfer PIPs and PC in vitro^22^. Therefore, we hypothesized that Sec14L6 could transfer lipids to promote LD biogenesis. First, we examined the lipid transfer activity of Sec14L6 using a FRET-based assay, as previously described^27^. Briefly, for measurement of PI4P transfer activities, donor liposomes containing DGS-NTA, PI4P, NBD-PH (PH domain of FAPP) and Rhodamine-PE (Rh-PE) were mixed with non-fluorescent ‘heavy’ (sucrose-loaded) acceptor liposomes containing DGS-NTA, PE, and DOPC, and measurements began after addition of purified Sec14L6 (Fig. 6A).

**Fig. 6.**
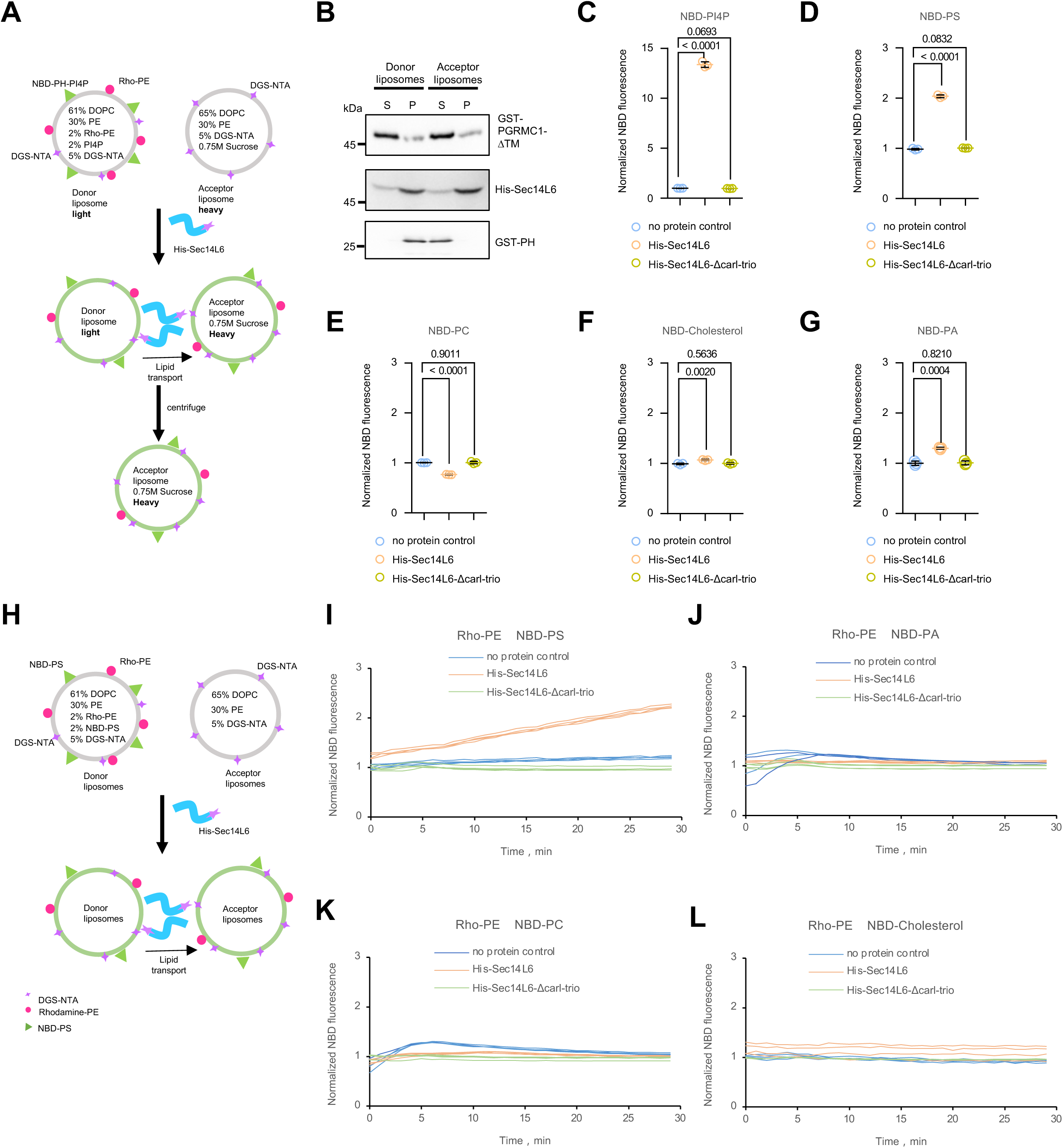
Sec14L6 transfers PI4P and PS in vitro. **A.** Schematic representation of the FRET-based *in vitro* lipid transfer assay of purified His-Sec14L6 using purified NBD-PH as a sensor for PI4P. **B.** Liposome pulldown assay confirmed the efficient binding of His-Sec14L6 and GST-PH to liposomes. GST-PGRMC1 proteins were used as a negative control. Western blots were performed with antibodies against GST and His. **C.** Donor liposomes (light; 20 μM) containing fluorescent lipids (2% PI4P, 2% Rhodamine-PE, 5% DGS-NTA, 61% DOPC, and 30% PE) with purified NBD-PH (250 nM) were mixed 1:1 with acceptor liposomes [heavy; 20 μM: 65% DOPC, 30% PE, and 5% DGS-NTA] in the absence or presence of His-Sec14L6 proteins (0.25 μM). The NBD fluorescence was normalized to the fluorescence intensity of no protein control. Ordinary one-way ANOVA with Tukey’s multiple comparisons test. Mean ± S.D. **D-G.** As in (**C**), donor liposomes (light; 20 μM) containing fluorescent lipids (2% NBD-PS (**D**), 2% NBD-PC (**E**), 2% NBD-Cholesterol (**F**) or 2% NBD-PA (**G**) with 2% Rhodamine-PE, 5% DGS-NTA, 61% DOPC, and 30% PE) were mixed 1:1 with acceptor liposomes [heavy; 20 μM: 65% DOPC, 30% PE, and 5% DGS-NTA] in the absence or presence of His-Sec14L6 (0.25 μM). Ordinary one-way ANOVA with Tukey’s multiple comparisons test. Mean ± S.D. **H**. Schematic representation of the FRET-based *in vitro* lipid transfer assay of purified His-Sec14L6 over time. **I-L.** Donor liposomes (20 μM) containing fluorescent lipids [2% NBD-PS (**I**), 2% NBD-PA (**J**), 2% NBD-PC (**K**), or 2% NBD-Cholesterol (**L**) with 2% Rhodamine-PE, 5% DGS-NTA, 61% DOPC, and 30% PE) were mixed 1:1 with acceptor liposomes [20 μM: 65% DOPC, 30% PE, and 5% DGS-NTA] in the absence or presence of His-Sec14L6 proteins (0.25 μM). The NBD fluorescence was normalized to the fluorescence intensity at time=0.

Liposome pelleting assays confirmed that His-Sec14L6 was associated with donor and acceptor liposomes via His-tag and Ni-NTA interaction, using GST-PGRMC1-ΔTM as a negative control (Fig. 6B). In addition, the purified GST-tagged PH domain of FAPP (GST-FAPP-PH) bound specifically to donor (with PI4P) but not to acceptor (without PI4P) liposomes (Fig. 6B). Furthermore, we examined the size distribution of liposomes by particle tracking analysis. Both donor and acceptor liposomes were well dispersed, uniform in size (∼100 nm), and stable in solution, suggesting that no spontaneous fusion or hemifusion of liposomes occurred (Fig. S5A).

Donor liposomes supplemented with PI4P were incubated with ‘heavy’ acceptor liposomes in the presence or absence of purified Sec14L6 protein, with purified His-Sec14L6-ΔCarl-Trio serving as a control. Donor and acceptor liposomes were then separated by centrifugation after stopping the reaction with a cocktail of imidazole and proteinase K. PI4P transfer from donor to acceptor liposomes was accompanied by an ∼8-fold increase in NBD fluorescence compared to the control (Fig. 6C), suggesting that Sec14L6 efficiently transfers PI4P *in vitro*.

Next, we examined the transfer activity of Sec14L6 for other glycerophospholipids and sterols. Interestingly, Sec14L6 could transfer PS (Fig. 6D), but not PC (Fig. 6E), PA (Fig. 6F) or cholesterol (Fig. 6G).

To further validate the result, we tracked the lipid transfer activity of Sec14L6 over time using fluorescence FRET-based assay as previously reported^36^ (Fig. 6H). We observed a slow but robust increase in NBD fluorescence over time after addition of His-Sec14L6 compared to the control (Fig. 6I), and the curve reached the plateau after about 90 minutes (Fig. S5B). However, we did not observe any significant increase in NBD fluorescence in donor liposomes containing NBD-PA (Fig. 6J), NBD-PC (Fig. 6K) or NBD-cholesterol and Rho-PE (Fig. 6L) in the presence of His-Sec14L6, indicating that Sec14L6 does not transfer PA, PC, PE, or cholesterol *in vitro*.

Moreover, the NBD fluorescence of the lipid transfer reactions as shown in Fig. 6H with or without His-Sec14L6 was almost the same after the further addition of dithionite (Fig. S5C), which quenched NBD fluorescence only in the outer leaflet of the liposome bilayer but did not affect the inner leaflet^37^, ruling out the possibility that the Sec14L6-mediated increase in fluorescence was due to fusion or hemifusion between donor and acceptor liposomes.

### The regulation of PS transfer activity of Sec14L6

Next, we investigated whether the PS transfer activity of Sec14L6 is subject to regulation. Interestingly, we found that the purified Carl-Trio domain, but not the NT or CT region, could transfer PS, which was more efficient than full-length Sec14L6 (Fig. 7A). Noteworthy, the purified Carl-Trio domain was able to mediate PS transfer between liposomes without DGS-NTA-Ni (Fig. 7B), consistent with the capacity of the Carl-Trio domain to target liposome membranes. These results suggested that the PS transfer activity of Sec14L6 is regulated by an auto-inhibition mechanism similar to LD association.

**Fig. 7.**
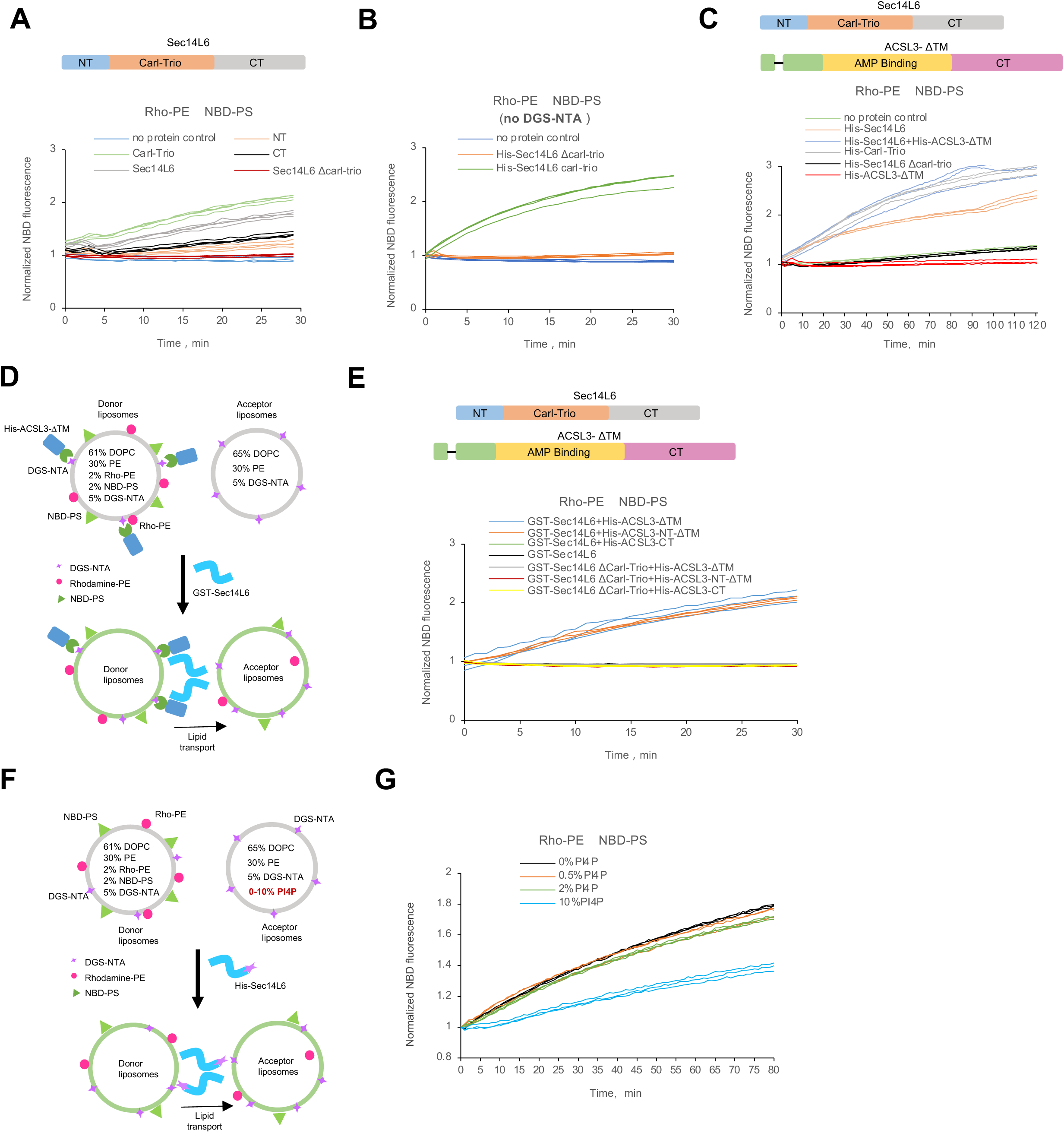
The regulation of the PS transfer activity of Sec14L6. **A.** As in Fig.6I, donor liposomes (20 μM) containing fluorescent lipids [2% NBD-PS, 2% Rhodamine-PE, 5% DGS-NTA, 61% DOPC, and 30% PE) were mixed 1:1 with acceptor liposomes [20 μM: 65% DOPC, 30% PE, and 5% DGS-NTA] in the absence (blue) or presence of either His-Sec14L6 (grey; 0.25 μM), His-Sec14L6-NT (orange; 0.25 μM), His-Sec14L6-CT (black; 0.25 μM) or His-Sec14L6-Carl-Trio (green; 0.25 μM). The NBD fluorescence was normalized to the fluorescence intensity at time=0. **B.** Donor liposomes (20 μM) containing fluorescent lipids [2% NBD-PS, 2% Rhodamine-PE, 66% DOPC, and 30% PE) were mixed 1:1 with acceptor liposomes [20 μM: 70% DOPC and 30% PE] with either His-Sec14L6-Carl-Trio (green; 0.25 μM) or His-Sec14L6ΔCarl-Trio (orange; 0.25 μM). No protein control was in blue. The NBD fluorescence was normalized to the fluorescence intensity at time=0. **C.** As in (**A**), *in vitro* PS transfer reactions in the absence (green) or presence of either His-Sec14L6 (orange; 0.25 μM), His-Sec14L6-Carl-Trio (grey; 0.25 μM) or the combination of His-Sec14L6 (blue; 0.25 μM) and His-ACSL3ΔTM (blue; 0.25 μM). His-Sec14L6ΔCarl-Trio (black; 0.25 μM) was used as negative control. **D.** Schematic representation of modified FRET-based in vitro lipid transfer assay of purified GST-Sec14L6 in presence of His-ACSL3 proteins. **E.** *in vitro* PS transfer reactions as in (**D**) showing the effects of different indicated His-ACSL3 proteins (0.25 μM for each) on the PS transfer activity of GST-Sec14L6 (0.25 μM) or GST-Sec14L6ΔCarl-Trio (0.25 μM). **F.** Schematic representation of modified FRET-based in vitro lipid transfer assay of purified His-Sec14L6 over time. **G.** Donor liposomes (20 μM) containing fluorescent lipids [2% NBD-PS, 2% Rhodamine-PE, 5% DGS-NTA, 61% DOPC, and 30% PE) were mixed 1:1 with acceptor liposomes (20 μM: 65% DOPC, 30% PE, and 5% DGS-NTA with 0, 0.5%, 2% or 10% PI4P) in the presence of His-Sec14L6 (0.25 μM). The NBD fluorescence was normalized to the fluorescence intensity at time=0.

We therefore hypothesized that the activity of Sec14L6 might be stimulated by ACSL3. Indeed, the addition of purified His-ACSL3-ΔTM increased the PS transfer activity of Sec14L6 to a level similar to that of the Carl-trio domain (Fig. 7C).

Remarkably, the addition of His-ACSL3-ΔTM or His-ACSL3-NT-ΔTM, but not the Sec14L6-binding defective mutant His-ACSL3-CT, stimulated the PS transfer activity of GST-Sec14L6 (Fig. 7D, E). This effect was mediated via the Carl-trio domain, as the Sec14L6 mutant lacking of this domain could not be stimulated by ACSL3 (Fig. 7E). Of note, GST-Sec14L6 was unable to transfer PS, likely due to its inability to target liposomes (Fig. 7E). These results suggest that ACSL3 enhances the PS transfer activity of Sec14L6 *in vitro*.

Since our results showed that Sec14L6 transferred PI4P and PS *in vitro*, we asked whether these two lipids competed with each other for binding to the same hydrophobic cavity in the Carl-Trio domain of Sec14L6. Indeed, the presence of PI4P in acceptor liposomes inhibited the PS transfer activity of Sec14L6 in a dose-dependent manner (Fig. 7F, G). Collectively, our results suggest that the PS transfer activity of Sec14L6 is regulated by ACSL3 and PI4P.

### Lipid transfer of Sec14L6 is required for LD biogenesis

Next, we asked whether the lipid transfer activity of Sec14L6 is required for LD biogenesis. Based on protein sequence alignment, we made four point mutants of Sec14L6 (space-filling mutants), in which conserved hydrophobic residues in the Carl-Trio domain were mutated to trptophan (79VV-80WW; 153F-W; 171L-W; 237L-W) (Fig. 8A, B). We measured the PS transfer activities of these space-filling mutants using the *in vitro* lipid transfer assay. These space-filling mutants caused a substantial decrease in PS transfer than WT Sec14L6, but they did not completely lose the transfer activities, as shown by a very small increase in NBD fluorescence (Fig. 8C).

**Fig. 8.**
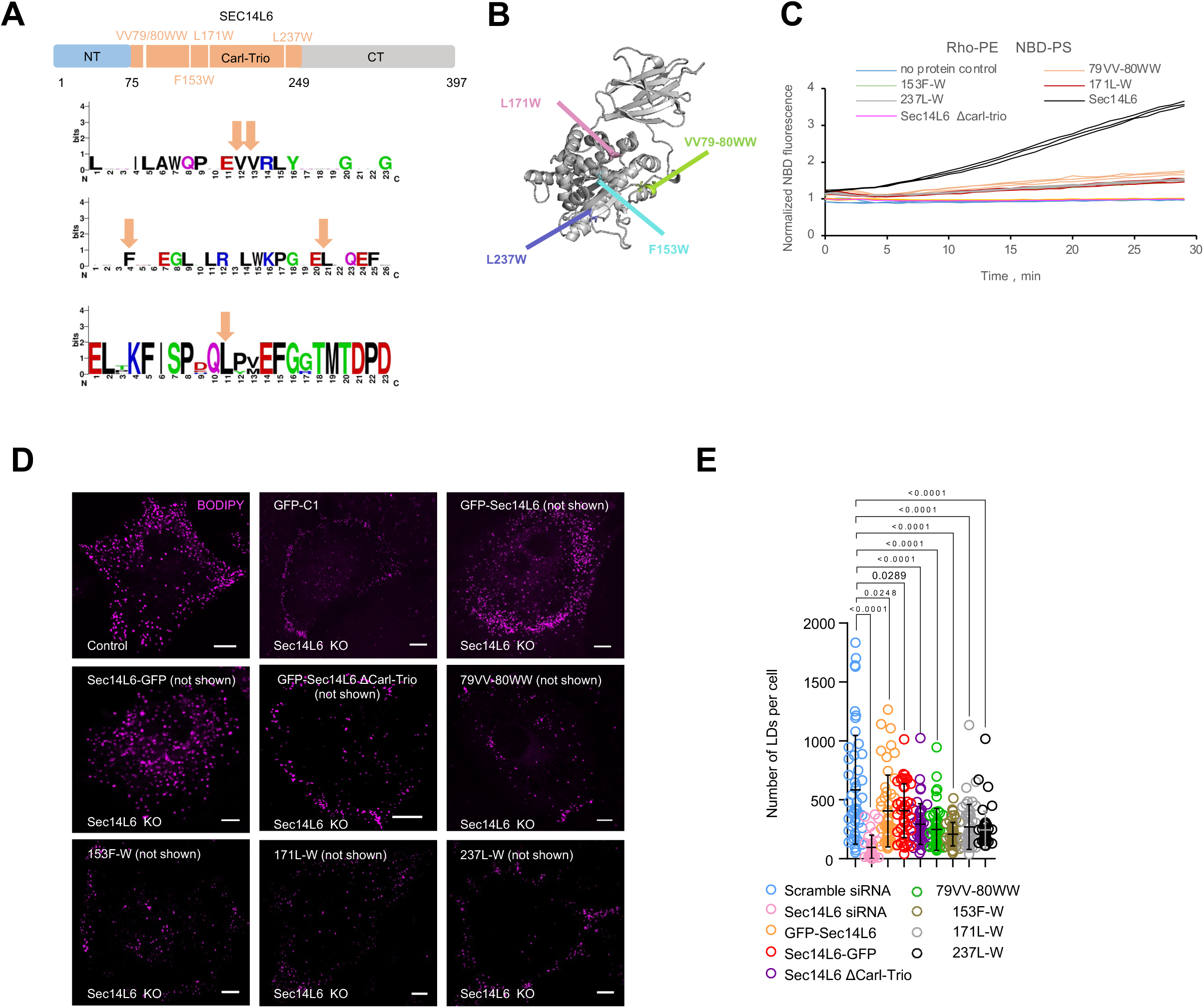
Lipid transfer activity of Sec14L6 is required for LD biogenesis. **A.** Schematic representation of conserved hydrophobic residues in the Carl-Trio domain of Sec14L6. **B.** AlphaFold-predicted structure showing conserved hydrophobic residues in the Carl-Trio domain of Sec14L6. **C.** Donor liposomes (20 μM) containing fluorescent lipids [2% NBD-PS, 2% Rhodamine-PE, 5% DGS-NTA, 61% DOPC, and 30% PE) were mixed 1:1 with acceptor liposomes [20 μM: 65% DOPC, 30% PE, and 5% DGS-NTA] in the absence (blue) or presence of either WT His-Sec14L6 (black; 0.25 μM), His-Sec14L6-VV79/80WW (orange; 0.25 μM), His-Sec14L6-F153W (green; 0.25 μM), His-Sec14L6-L171W (red; 0.25 μM) or His-Sec14L6-L237W (grey; 0.25 μM). The NBD fluorescence was normalized to the fluorescence intensity at time=0. **D.** Representative images of live BODIPY (magenta)-stained control HeLa or Sec14L6 KO#2 clones rescued with GFP empty vector, WT GFP-Sec14L6, Sec14L6-GFP, GFP-Sec14L6-ΔCarl-Trio, GFP-Sec14L6-VV79/80WW, GFP-Sec14L6-F153W, GFP-Sec14L6-L171W or GFP-Sec14L6-L237W. **E.** The number of LDs in cells as shown in (**D**) in more than 3 independent experiments. Ordinary one-way ANOVA with Tukey’s multiple comparisons test. Mean ± SD. Scale bar, 10μm in (D).

Furthermore, we investigated whether and to what extent LD formation is dependent on the lipid transfer activity of Sec14L6. Live-cell imaging revealed that the defect in LD formation was specific to Sec14L6, as the introduction of WT Sec14L6 significantly rescued the phenotype (Fig. 8D). Importantly, these four mutants with PS transfer deficiencies did not significantly rescue the defect in LD number caused by Sec14L6 KO (Fig. 8E). These results suggest that the PS transfer activity of Sec14L6 is required for the formation of LDs.

### Sec14L6 promotes the differentiation of adipose-derived mesenchymal stem cells

We next investigated the potential physiological functions of Sec14L6 in human adipose-derived mesenchymal stem cells (ADSCs)^38^ that were able to differentiate into adipocytes after stimulation with the formation of numerous LDs as hallmarks of mature adipocytes (Fig. 9A). In the control, we observed that the number or size of LDs increased greatly after induction (up to 16 days) (Fig. 9B, H). Of note, we found that some giant LDs that were clearly seen in bright-field (BF) images at the late time points (>10 days), were not efficiently stained with BODIPY (Fig. 9B-G). Therefore, we quantified the size or number of LDs using BODIPY and BF images. Suppression of Seipin or FTO, two essential factors for ADSC differentiation^9, 39^, almost completely blocked the formation of LDs (Fig. 9C, D, H; Fig. S5D, E). Importantly, suppression of Sec14L6 by siRNAs strongly impaired LD formation during differentiation after induction (Fig. 9E, H; Fig. S5F), to a similar extent as ACSL3 (Fig. 9F, H; Fig. S5G), but not as strongly as Seipin or FTO. Interestingly, PGRMC1 (Fig. 9G; Fig. S5H), but not PGRMC2 (not shown), significantly reduced LD formation upon induction, supporting a role of PGRMC1 in LD biogenesis.

**Fig. 9.**
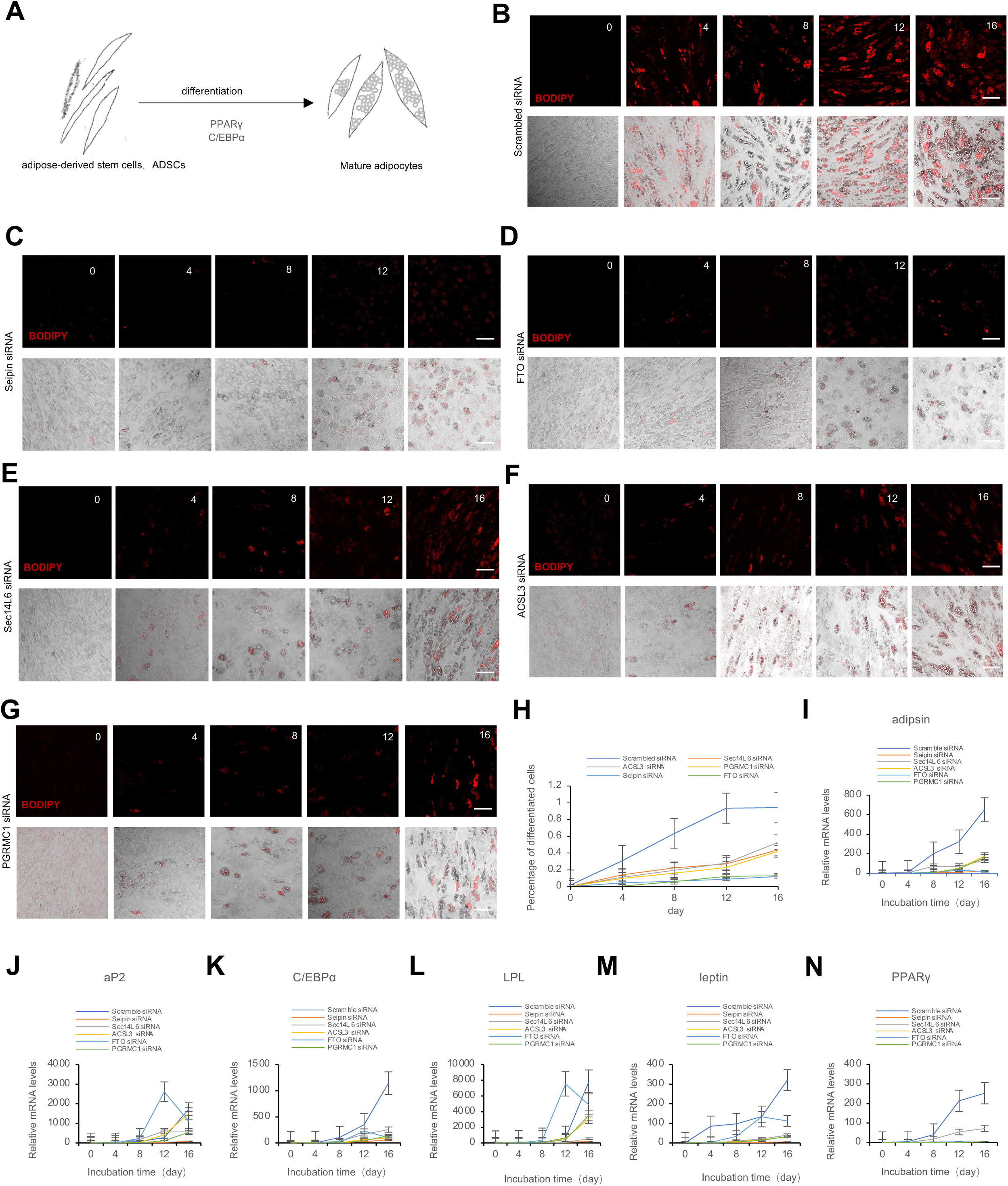
A role of Sec14L6 in differentiation of adipose stem cell differentiation. **A.** Schematic representation of differentiation of adipose-derived stem cells (ADSCs) with the formation of numerous LDs as one of hallmarks in mature adipocytes. **B-G**. Representative images of live BODIPY (red)-stained ADSCs at time=0, 4, 8, 12, 16 day, treated with scrambled (**B**), Seipin (**C**), FTO (**D**), Sec14L6 (**E**), ACSL3 (**F**) or PGRMC1 (**G**) siRNAs. Top panel: BODIPY channel; bottom panel: merged images of BODIPY and DIC channels. **H.** Percentage of mature ADSCs in (**B-G**) in more than 3 independent experiments. Mean ± SD. Mature ADSCs were defined by the presence of visibly large LDs marked by BODIPY and were further confirmed in bright field images. **I-N**. qPCR showed the mRNA levels of factors in ADSC differentiation, including adipsin (**I**), aP2 (**J**), C/EBPα (**K**), LPL (**L**), leptin (**M**) or PPARγ (**N**) as in (**I-N**), in more than 3 independent experiments. Mean ± SD. Scale bar, 50μm in (B-G).

Next, we confirmed the role of Sec14L6 in ADSC differentiation by examining adipocyte-specific factors, such as adipsin^40^, aP2^41^, leptin^42^, C-EBPα and PPARγ^43^. Accordingly, Seipin, FTO, or ACSL3 are essential for adipocyte differentiation, as their suppression led to in a significant reduction in transcriptional levels of these factors (Fig. 9I-N). Importantly, the mRNA levels of these factors were significantly reduced in Sec14L6- or PGRMC1-depleted cells (Fig. 9I-N), suggesting an important role of Sec14L6 and PGRMC1 in this process. Taken together, our results demonstrate that Sec14L6 and its interactors ACSL3 and PGRMC1 are required for LD formation and involved in ADSC differentiation.

## Discussion

In this study, we identified Sec14L6 as a PS/PI4P transporter indispensable for LD formation and differentiation of ADSCs. Sec14L6 directly interacted with ACSL3. The interaction facilitated the association of Sec14L6 with LDs, and stimulated the PS transfer activity of Sec14L6 *in vitro*. We further identified PGRMC1 as an adaptor that recruits Sec14L6 to the ER (Fig. S5I). We propose that Sec14L6 transfers PS and/or PI4P between the ER and LDs to promote LD formation (Fig. S5J).

A recent study showed that ORP5 cooperates with ORP8 to regulate LD biogenesis and maintenance at ER-mitochondrial MCS^28^. In addition, Orp5 was shown to regulate lipid homeostasis of LD membranes at ER-LD interfaces^29^. Loss of ORP5 resulted in enlarged LDs with PS deficiency in LD membranes. While both Sec14L6 and Orp5 are PS/PI4P transporters involved in lipid exchange between the ER and LDs, they differ in other aspects. First, the ORP5 gene is widely distributed in metazoans. In contrast, Sec14L6 is only found in large mammals. For example, this gene is not present in the mouse. Second, ORP5 is a tail-anchor protein that is anchored to ER membrane via a TM domain and can be recruited to ER-plasm membrane (PM)^44^ and ER-LD membrane contact sites (MCS)^29^, where it functions as a tether. However, Sec14L6 is mainly cytosolic and not membrane-bound. The association of Sec14L6 with the ER or LDs is highly regulated and depends on its interactors. More importantly, Sec14L6 did not function as a tether at ER-LD MCS even in the presence of ACSL3-Halo and PGRMC1-mCh (unpublished results). Therefore, we believe that Sec14L6 is a cytosolic lipid transporter that transfers lipids in a different way from ORP5. Although both Sec14L6 and ORP5 can transfer PS and PI4P, their expression specificity, localization, and mode of lipid-transfer were clearly different from each other, which may account for the different phenotypes in loss-of-function experiments.

In addition to ORP5, other LTPs have also been localized to ER–LD interfaces, including ORP2^16^, VPS13A and VPS13C^17, 18^, and ATG2^45^. Why are these LTPs present at ER-LD interfaces? One plausible explanation is that these LTPs transfer various lipids, e.g., PS, PC, PE or cholesterol, from the ER to LDs in different functional contexts, including the formation of LDs in the ER, the growth of mature LDs out of the ER, the maintenance of lipid homeostasis of LDs and adaptation to stress. Further work is required to investigate the spatial and functional relationships between Sec14L6, ORP5, VPS13 proteins and ATG2.

A unique property of PS is its negative charge, which allows PS to control the localization and activity of a number of proteins, including kinases and phospholipases^46, 47^. In addition, PS is important for the organization of LD membranes^29^. Given that the amount of PS on the LD surface is very small (<1% of total phospholipids)^48^, PS probably plays an important signaling/regulatory role during the formation of nascent LDs, but not a structural role. Indeed, Guyrad et al. proposed that Orp5 might create a local phospholipid environment in the ER important for the proper action of the LD assembly complex^28^, including Seipin, which binds negatively charged phospholipids, such as PS. Therefore, it is also plausible that Sec14L6 regulates local ER phospholipids important for LD assembly complexes.

Defects in LD biogenesis are the hallmark of numerous metabolic and non-metabolic diseases such as type II diabetes, cancer, heart disease or viral infections^49, 50^. Therefore, our results will provide new mechanistic insights for these disorders. Remarkably, Sec14L6 is only found in large mammals, our study therefore reveals a unique regulator of LD biogenesis that is specific to large mammals.

## Method

### Cell culture, transfection, RNAi

Human cervical cancer HeLa cells (ATCC), and human embryonic kidney 293T (ATCC) were grown in (Invitrogen) supplemented with 10% fetal bovine serum (Gibco). All of the cell lines used in this study were confirmed free of mycoplasma contamination.

Transfection of plasmids and RNAi oligos was carried out with Lipofectamine 2000 and RNAi MAX, respectively. For transfection, cells were seeded at 4 x 10^5^ cells per well in a six-well dish ∼16 h before transfection. Plasmid transfections were performed in OPTI-MEM (Invitrogen) with 2 μl Lipofectamine 2000 per well for 6 h, followed by trypsinization and replating onto glass-bottom confocal dishes at ∼3.5 x 105 cells per well. Cells were imaged in live-cell medium (DMEM with 10% FBS and 20 mM Hepes with no penicillin or streptomycin) ∼16–24 h after transfection. For siRNA transfections, cells were plated on 3.5 cm dishes at 30–40% density, and 2 μl Lipofectamine RNAimax (Invitrogen) and 50 ng siRNA were used per well. At 48 h after transfection, a second round of transfection was performed with 50 ng siRNAs. Cells were analyzed 24 h after the second transfection for suppression.

### CRISPR-Cas9-mediated gene editing

To make Sec14L6 KO HeLa cell lines, two gRNAs (5,-CTCCTGTGTATGCATCCAGG-3, and 5,-AACGGCATATGCGGCCACGA-3,) were used to delete ∼38 bp from exon 23 of Sec14L6 gene .Complementary gRNAs were annealed and subcloned into the pSpCas9(BB)-2A-GFP (pX-458) vector (Addgene 48138) using BbsI. Upon transfection, HeLa cells were grown in an antibiotic-free medium for 48 h, followed by single-cell sorting by fluorescence-based flow cytometry.

To make SEC14L6-GFP-KI HeLa cell lines, a single gRNA (5’-GAAATTCTAGGTGAACCTCA-3,) were used to target the N-terminus of the Sec14L6 gene (Fig. S3C). HeLa cells were transfected with plasmids encoding the gRNA and a donor construct containing sfGFP and two homologous arms using Lipofectamine 2000. 48 h after transfection, single clones were sorted by flow cytometry. A positive clone was verified by DNA gels (Fig. S3D) and immunoblots (Fig. S3E).

### Plasmids

Sec14L6(NM_001193336), ACSL3(NM_203372.1), PGRMC1(NM_006667) were cloned from HeLa cDNA library. The coding sequence (CDS) of Sec14L6 and its truncation mutants were cloned into mEGFP-C1(addgene 54579) between SalI and SacII. 14xHis-NEDD8-Sec14L6 was constructed by cloning its CDS to 14xHis-NEDD8 vector. The CDS of ACSL3, PGRMC1 and their truncation mutants were cloned into Halo-N1 (addgene 54767) using SacII and EcoRI. GST-Sec14L6 was constructed by cloning the Sec14L6 CDS into PGEX-2T vector using BamHI and EcoRI. 14xHis-NEDD8-PGRMC1 and 14xHis-NEDD8-ACSL3 were constructed by cloning the ORFs into 14xHis-NEDD8 vector using BamHI and HindIII. The FAPP-PH construct was a gift from Dr. Shunji Jia (Institute of Genetics and Developmental Biology, Chinese Academy of Sciences).

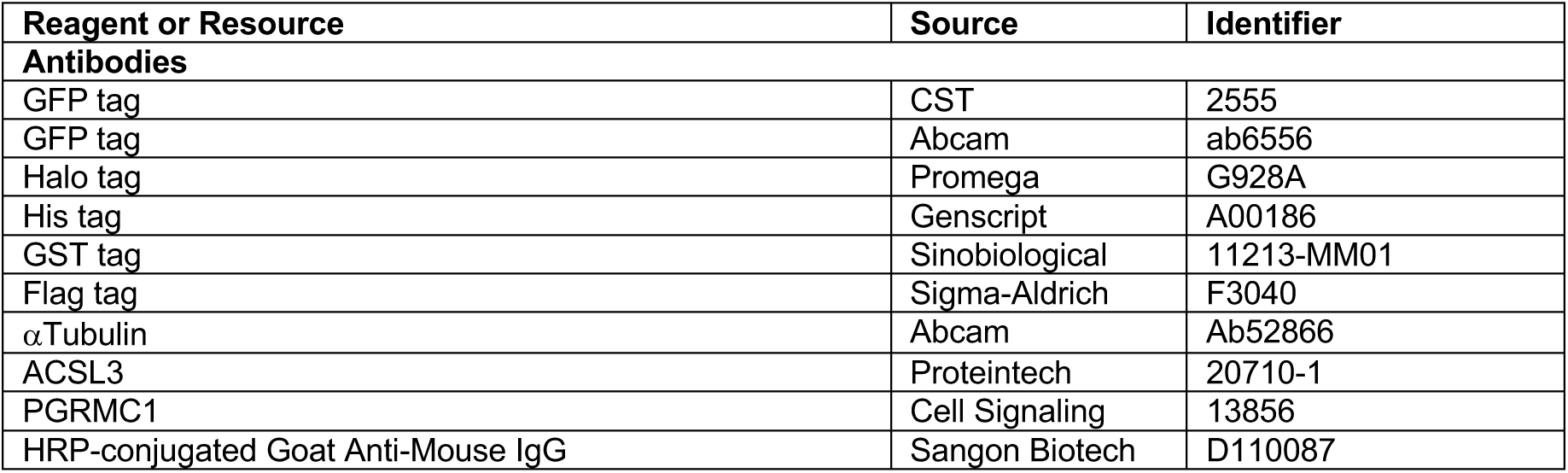
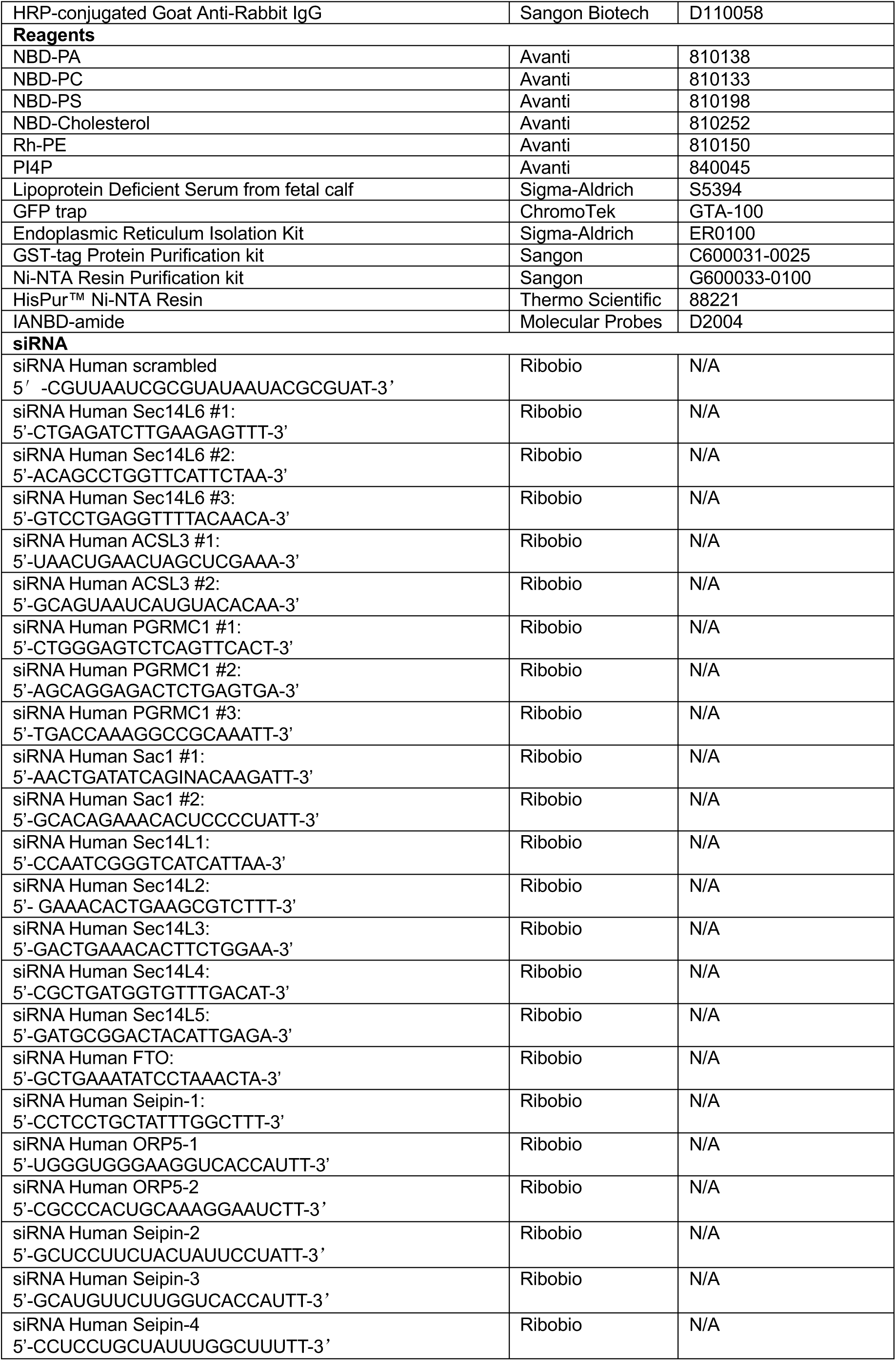
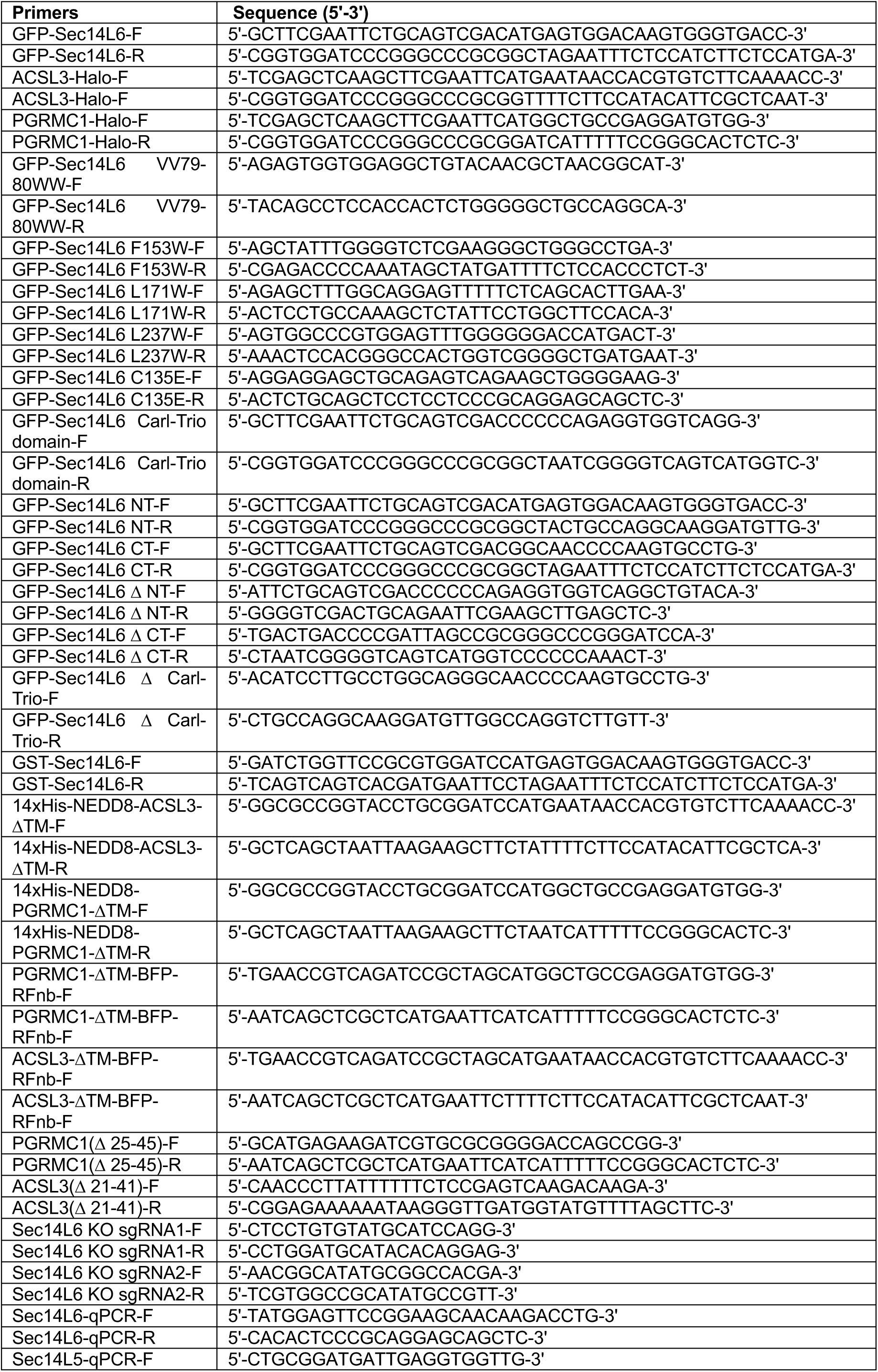
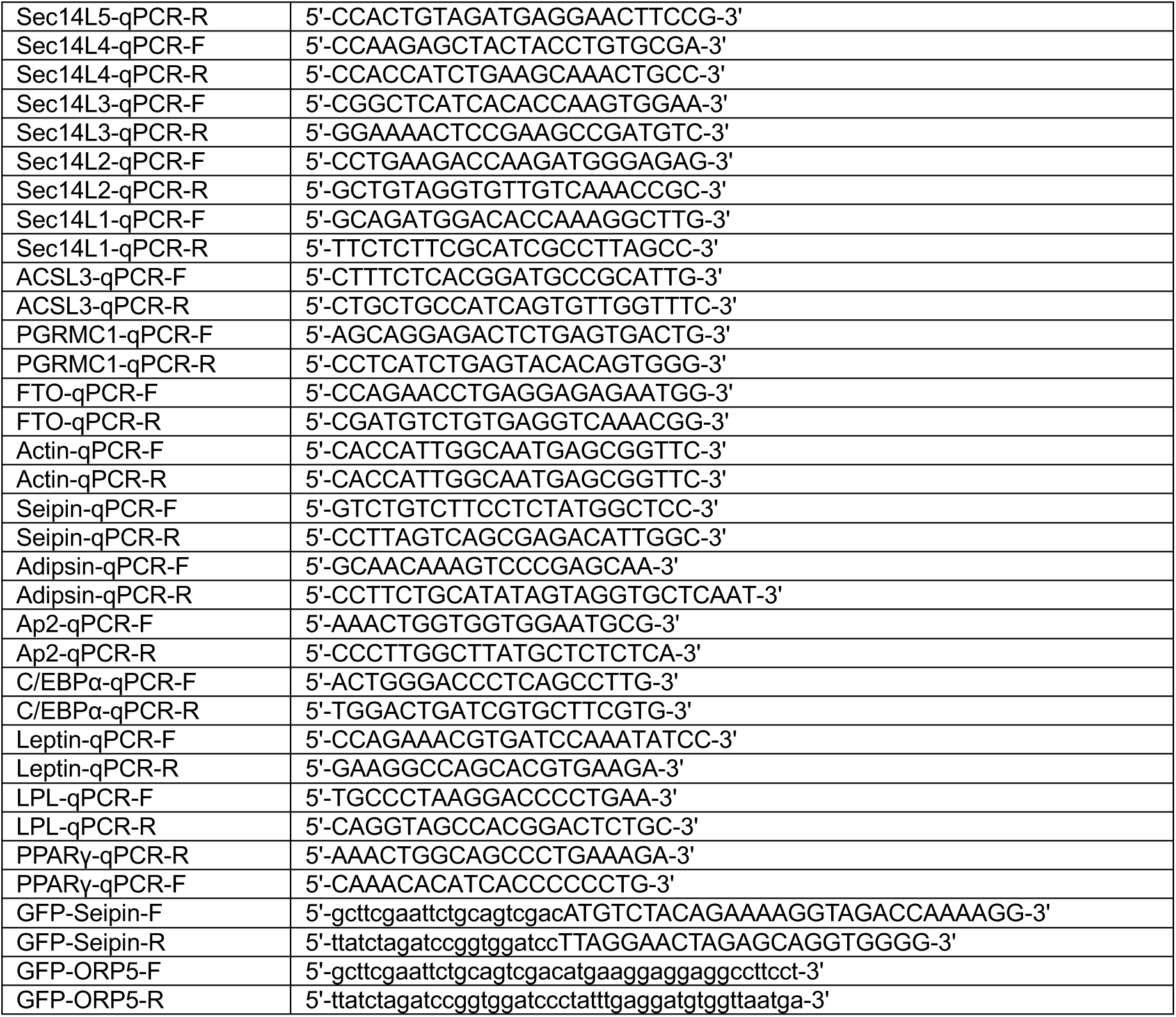

### Western Blotting

After being heated to 95°C for 10 min, the SDS sample buffer was used to lyse the cells. The resulting samples were subjected to SDS-PAGE electrophoresis and transferred to a NC Transfer Membrane (HATF00010; Millipore). The membrane was then blocked with 5% (m/v) non-fat powdered milk at room temperature for 1 h and then incubated with primary antibodies at 4°C overnight. After washing three times, membranes were incubated with secondary antibodies that were conjugated with horseradish peroxidase (1/10,000) for 1 h at room temperature. Visualization was performed using enhanced chemiluminescence (P0018M-2; Beyotime)

### Purification of ER membranes by density gradient centrifugation

ER fractions were enriched using Endoplasmic Reticulum Isolation Kit (ER0100; Sigma-Aldrich) according to the manufacturer’s instructions. Briefly, scrambled or PGRMC1 siRNA-treated HeLa cells stably expressing GFP empty vector or GFP-Sec14L6 from five confluent 10-cm dishes were collected, followed by centrifugation at 600 × g for 5 min. After washing the cells three times with PBS, the packed cell volume (PCV) was measured and then suspended in a volume of hypotonic extraction buffer (10 mM HEPES, pH 7.8, with 1 mM EGTA and 25 mM potassium chloride) equivalent to three times the PCV. After the incubation of the cells for 20 min at 4°C allowing the cells to swell, the cells were centrifuged at 600 × g for 5 min, followed by the measurement of the “new” PCV. After adding a volume of isotonic extraction buffer (10 mM HEPES, pH 7.8, with 0.25 M sucrose, 1 mM EGTA, and 25 mM potassium chloride) equivalent to two times the “new” PCV, the suspension was then transferred to a 7-ml Dounce homogenizer, followed by the lysis of the cells with 10 strokes and then the centrifugation of the homogenate at 1,000 × g for 10 min at 4°C. After the transfer of the supernatant to another centrifuge tube, the supernatant was centrifuged at 12,000 × g for 15 min at 4°C, followed by another centrifugation for 60 min at 100,000 × g at 4°C. The pellet was the microsomal fraction and further verified by Western blots using anti-Calnexin antibody.

### Isolation of LD fractions by density gradient centrifugation

Isolation of LD fractions from HeLa cells or Sec14L6-GFP-KI HeLa line was performed as previously reported^51^ with some modifications. Briefly, cells were washed and lysed on ice using a Potter-Elvehjem tissue homogenizer in Hypotonic lysis medium (HLM, 20 mM Tris-Cl, pH 7.4; 1 mM EDTA; Protease inhibitors). The cell lysates were centrifuged at 1,000 g for 10 min at 4°C, followed by addition of 1/3 volume of ice-cold HLM containing 60% sucrose (final 20% sucrose) into the supernatant and the floating fat layer. After aggregates of LDs were finely and thoroughly dispersed, the sample was layered into the bottom of a 13.2-ml ultracentrifuge tube for an SW41Ti rotor. 5 mL ice-cold HLM containing 5% sucrose and 5 mL ice-cold HLM were sequentially layered over the sample, followed by centrifuge at 28,000 g for 30 min at 4°C. The LD fraction was then transferred to a microcentrifuge tube, and was centrifuged at ∼20,000 g for 10 min at 4°C.The LD fraction was resuspended in ice-cold HLM, followed by checking purity of the LD fraction by SDS-PAGE of solubilized proteins and immunoblotting.

### De-lipidation and LD formation induction

Cell de-lipidation and LD formation induction were conducted as described previously^10^. Briefly, cells were delipidated by culturing in serum-free medium supplemented with 5% lipoprotein-deficient serum (Sigma) for 60 h. LD formation was induced by adding 0.2 mM OA (final concentration) for the indicated times.

### TAG measurement

The indicated HeLa cells cultured in 6-well plates were lysed in 1% Triton X-100, and cell extracts were used to measure the TAG content by Triglyceride Colorimetric Assay Kit based on glycerol phosphate oxidase-based assays (GPO-PAP method) (BioSino Bio) according to the manufacturer’s instructions. Briefly, the total protein concentration of each sample was determined by BCA assay (Thermo Fisher), and reactions were monitored in microplate reader at optical density 510 nm.

### GFP-trap assay

GFP trap (GTA-100; ChromoTek) was used for the detection of protein–protein interactions, and the GFP-Trap assays were performed according to the manufacturer’s protocol. Briefly, after 24 h transfection with the indicated plasmids, cells were lysed in ice-cold lysis buffer (50 mM Tris-HCl, pH 7.5, 150 mM NaCl, 1 mM EDTA, 1% Triton X-100 and protease inhibitor cocktail). Lysates were centrifuged at 13,000 rpm for 10 min at 4°C and pellets were removed. Supernatants were incubated with GFP-Trap agarose beads for 1 h at 4°C with gentle rocking. After washing four times with lysis buffer, beads were boiled with SDS sample buffer. Proteins of interest were analyzed by immunoblotting. 5% input was used in GFP traps unless otherwise indicated.

### GST tag and his tag protein purification

GST and His constructs were transformed into Escherichia coli BL21 (DE3) cells, and cells were incubated at 37°C until the optical density (OD) at 600 nm reached 0.6–0.8. Subsequently, cells were incubated at 16°C for another hour, followed by induction with 1 mM IPTG overnight at 16°C.Cells were lysed via sonication. GST fusion proteins were purified via the GST-tag Protein Purification kit (C600031-0025, Sangon, China), His fusion proteins were purified via the Ni-NTA Sefinose (TM) Resin Purification kit (G600033-0100, Sangon, China)

### *In vitro* pull-down assays of GFP tag and his tag

HEK293 cells transiently transfected with GFP-Sec14L6 (full length and truncations) were lysed in high-salt lysis buffer (RIPA buffer containing 500 mM NaCl, proteasome inhibitors and PMSF). GFP-Trap beads were used to pellet GFP-Sec14L6 (full-length and truncations) from cell lysates, followed by washing with high-salt lysis buffer for 10 times. The GFP beads were incubated with Purified His-ACSL3 or His-PGRMC1 overnight at 4°C, respectively, followed by washing beads with freshly prepared HNM buffer (20 mM Hepes, pH 7.4, 0.1 M NaCl, 5 mM MgCl2, 1 mM DTT and 0.2% NP-40). GFP beads were resuspended in 100 μL 2 x SDS-sampling buffer. Re-suspended beads were boiled for 10 min at 95°C to dissociate protein complexes from beads. Western blotting was performed using anti-GFP, ACSL3 or PGRMC1 antibodies. The Coomassie staining was performed for purified His-ACSL3 and His-PGRMC1.

### Live imaging by high-resolution confocal microscopy

Cells were grown on 35 mm glass-bottom confocal MatTek dishes, and the dishes were loaded to a laser scanning confocal microscope (LSM900, Zeiss, Germany) equipped with multiple excitation lasers (405 nm, 458 nm, 488 nm, 514 nm, 561 nm and 633 nm) and a spectral fluorescence GaAsP array detector. Cells were imaged with the 63×1.4 NA iPlan-Apochromat 63 x oil objective using the 405 nm laser for BFP, 488 nm for GFP, 561nm for OFP, tagRFP or mCherry and 633nm for Janilia Fluo® 646 HaloTag® Ligand.

### Measurements of LD number and size

The LD number per HeLa or ADSC was counted manually with the help of an plugin of ImageJ (1.54i; NIH), Cell Counter. The LD size was quantified by manually measuring the diameter of individual LD by ImageJ. In case of clustered LDs in Sec14L6 KO cells, the area of individual LD was manually measured by ImageJ based on SEM micrographs.

### FRET

HeLa cells transiently transfected with Clover-Sec14L6-mRuby2 and ACSL3-Halo were imaged using a Dragonfly confocal microscopy system (Andor) equipped with a 60 × N.A. 1.4 oil-immersion objective. Clover, mRuby2, and FRET fluorescence was collected with the following parameters: Clover: 488 nm excitation, 521 ± 19 nm emission; mRuby2:561 nm excitation, 594 ± 21.5 nm emission; FRET: 488 nm excitation, 617 ± 39.5 nm emission. Image analysis was performed with Matlab7.0 software to calculate *N*_FRET_ (Normalized FRET) according to the equation as previously described^52^: 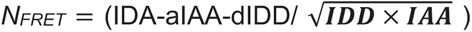, where I_DA_, I_DD,_ and I_AA_ are the background-subtracted FRET, Clover, and mRuby2 images, respectively.

### Mass spectrometry for identification of GFP-Sec14L6-interacting Proteins

The identification of Sec14L6-interacting proteins by MS was described in our previous study^53^. Briefly, the bound proteins were extracted from GFP-Trap agarose beads using SDT lysis buffer (4% SDS, 100 mM DTT, 100 mM Tris-HCl, pH 8.0), followed by sample boiling for 3 min and further ultrasonicated. Undissolved beads were removed by centrifugation at 16,000 g for 15 min. The supernatant, containing proteins, was collected. Protein digestion was performed with the FASP method. Briefly, the detergent, DTT, and IAA in the UA buffer were added to block-reduced cysteine. Finally, the protein suspension was digested with 2 µg trypsin (Promega) overnight at 37°C. The peptide was collected by centrifugation at 16,000 g for 15 min. The peptide was desalted with C18 StageTip for further LC-MS analysis. LC-MS/MS experiments were performed on a Q Exactive Plus mass spectrometer that was coupled to an Easy nLC (Thermo Fisher Scientific). The peptide was first loaded to a trap column (100 µm × 20 mm,5 µm, C18, Dr. Maisch GmbH) in buffer A (0.1% formic acid in water). Reverse-phase high-performance liquid chromatography (RP-HPLC) separation was performed using a self-packed column (75 µm × 150 mm; 3 µm ReproSil-Pur C18 beads, 120 °A; Dr. Maisch GmbH, Ammerbuch) at a flow rate of 300 nl/min. The RP-HPLC mobile phase A was 0.1% formic acid in water and B was 0.1% formic acid in 95% acetonitrile. The gradient was set as follows: 2–4% buffer B from 0 to 2 min, 4–30% buffer B from 2 to 47 min, 30–45% buffer B from 47 to 52 min, 45–90% buffer B from 52 min and to 54 min, and 90% buffer B kept until to 60 min. MS data were acquired using a data-dependent top20 method dynamically choosing the most abundant precursor ions from the survey scan (350–1,800 m/z) for HCD fragmentation. A lock mass of 445.120025 Da was used as the internal standard for mass calibration. The full MS scans were acquired at a resolution of 70,000 at m/z 200, and 17,500 at m/z 200. The maximum injection time was set to 50 ms for MS and 50 ms for MS/MS. Normalized collision energy was 27 and the isolation window was set to 1.6 Th. Dynamic exclusion duration was 60 s. The MS data were analyzed using Max Quant software version 1.6.1.0. MS data were searched against the UniProtKB human database (36,080 total entries, downloaded 2019.06.25). Trypsin was selected as the digestion enzyme. A maximum of two missed cleavage sites and a mass tolerance of 4.5 ppm for precursor ions and 20 ppm for fragment ions were defined for database search. Carbamidomethylation of cysteines was defined as a fixed modification, while acetylation of protein N-terminal and the oxidation of methionine were set as variable modifications for database searching. The database search results were filtered and exported with a <1% false discovery rate (FDR) at peptide-spectrum-matched level and protein level, respectively.

### Protein expression and purification

The His-Sec14L6 constructs were transfected into 30x 10-cm dishes of HEK293T cells, and the medium was changed 5 h later. After another 24 h, cells were incubated in OptiMEM. The medium was collected 48 h later, and was centrifuged at 500 g for 10 min. The supernatant was mixed with 500 µl of balanced HisPur™ Ni-NTA Resin (88221, Thermo Scientific) and incubated in a 4°C shaking bed for 2 h. After centrifugation at 1000 g for 10 min, the supernatant was discarded. Proteins on beads were washed with 20 mM Tris-HCl pH 8.0, 300 mM NaCl buffer containing 20 mM, 50 mM, 100 mM, and 200 mM imidazole sequentially. The eluted proteins were examined by Coomassie staining and western blotting. Then, the eluted proteins were concentrated (Millipore UFC900396), and washed with imidazole-free buffer to obtain the imidazole-free and concentrated His-Sec14L6 proteins.

### Liposome pelleting assay

For liposome pelleting assays, proteins were diluted to ∼7 µg/ml in osmotically matched protein dilution/HK buffer (20 mM Hepes, 120 mM NaCl, 1 mM EGTA, 0.2 mM CaCl_2_, 1.5 mM MgCl_2_, 1 mM DTT, 5 mM KCl, pH 7.4, 1% BSA) and were pre-cleared by ultracentrifugation at 120,000 g for 45 min using a TLA100.3 rotor in a Beckmann Optima-MAX benchtop ultracentrifuge. Heavy liposomes were prepared in the HK buffer with 0.75 M sucrose. 1 mL of pre-cleared protein solution was then mixed with 10 µL of the heavy liposomes and incubated with shaking at 25°C for 15 min. Liposomes were recovered by ultracentrifugation (16,100 g for 15 min), and supernatant and pellet fractions were resuspended in equal volumes of 1× Laemmli buffer and analyzed by western blots.

### *In vitro* FRET-based lipid transfer assay

Lipid transfer assays shown in Fig. 6A-G were performed as follows. Light liposomes (DOPC: liver PE: PI4P: Rhodamine-PE: DGS-NTA-Ni = 61: 30: 2: 2: 5) were prepared in the HK buffer and coated with 500 μM NBD-PH protein. Heavy liposomes (DOPC: liver PE: DGS-NTA-Ni = 65: 30: 5, 0.75 M sucrose) were prepared in the HK buffer. The transfer reaction was initiated by mixing two liposome populations, with or without 0.25 μM Sec14L6 proteins. After 15-min incubation at 25 °C, a cocktail of imidazole and proteinase K was added to terminate the transfer process, and the liposome suspensions were separated by centrifugation at 16,100 g for 15 min. Heavy liposomes were collected and resuspend with 1 mL HK buffer, followed by centrifuge at 16,100 g for 15min. This step was repeated for three times. After three additional washes with HK buffer, heavy liposomes were collected for NBD signal measure at 528 nm after excitation at 460 nm at 30°C using a CLARIOstar Plus Microplate Reader (BMG LABTECH).

Lipid transfer assays shown in Fig. 6H-L and Fig. 7&8 were conducted according to the protocols described in previous studies^27, 36^. Briefly, lipid transfer reactions were performed in 100 µL volumes in 96-well plates containing a protein: lipid ratio of 1:800, with 0.25 μM proteins, 20 μM donor liposomes (61% DOPC, 30% liver PE, 2% NBD-PA/NBD-PS/NBD-PC/NBD-Cholesterol, 2% Rhodamine-PE, and 5% DGS-NTA-Ni) and 20 μM acceptor liposomes (65% DOPC, 30% liver PE and 5% DGS-NTA-Ni). The fluorescence intensity of NBD was measured via excitation at 460 nm and detection at 538 nm every 1 min for indicated time at 30°C using a CLARIOstar Plus Microplate Reader. All data were normalized to the NBD fluorescence at first time point (t = 0).

### Dithionite assay

After performing the lipid transfer reaction as shown Fig. 6H, 2.5 μL freshly prepared dithionite buffer (100 mM dithionite in 50 mM Tris-HCl, pH=10) was added to reactions, and NBD fluorescence was monitored for an additional 15 min.

### Nanoparticle tracking analysis

The size distribution of liposomes was assessed by nanoparticle tracking analyses according to manufacturer instructions (NanoSight NS300, Malvern Panalytical). Briefly, the total number of particles was measured in light scatter mode without the use of an optical filter. Samples were applied with a constant flow rate using the supplied syringe-pump. liposomes were diluted 1:100 in HK buffer (filtered with a 0.2 μm filter) prior to NTA measurements. Three to five videos of 60 s were recorded per sample with a sCMOS camera at 25 frames per second (1,498 frames per video) and data were analyzed with the NTA software 3.1 (Build 3.1.54).

### NBD-FAPP-PH

NBD-PH-FAPP molecular probes was prepared as previously described^27^. Briefly, after removing dithiothreitol, purified PH-FAPP was mixed with a tenfold molar excess of N,N’-dimethyl-N-(iodoacetyl)-N’-(7-nitrobenz-2-oxa-1,3-diazol-4-yl) ethylenediamine (IANBD-amide, Molecular Probes). The labelling reaction was conducted overnight at 4 °C and terminated by adding a tenfold molar excess of l-cysteine. The free probe was removed by gel filtration and the labelled protein was analyzed by UV-visible spectroscopy to estimate labelling efficiency (∼100%), which was calculated from the ratio of the optical density (OD) of tyrosine and tryptophan at 280 nm (ε = 29,450 M cm^−1^ for PH-FAPP) and NBD at 495 nm (ε = 25,000 M cm^−1^).

### Differentiation of human adipose derived mesenchymal stem cells

Human adipose derived mesenchymal stem cells were cultured to confluence before differentiation in a 24-well plate. For adipogenesis, the cells cultured in DMEM with 10 % FBS were stimulated with a differentiation cocktail (Pricella PD-006, ProCell), which included 1-methyl-3 isobutylxanthine (IBMX), dexamethasone, human insulin and rosiglitazone in addition to pennicillin-streptomycin, and glutamine (differentiation medium) for 3 days. Then the cells were cultured in medium containing a maintaining cocktail (Pricella PD-006, ProCell) including glutamine, pennicillin-streptomycin and human insulin (maintaining medium) for 1 day and then cells were changed back to the differentiation medium. Cells were cultured in the two types of media alternately until sufficient LDs are visible in the stem cells. Mature ADSCs were defined by the presence of visibly large LDs marked by BODIPY and were further confirmed in bright field images.

### Statistical analysis

All statistical analyses and p-value determinations were performed in GraphPad Prism8.0.1. All the error bars represent Mean ± SD. To determine p-values, ordinary one-way ANOVA with Tukey’s multiple comparisons test was performed among multiple groups and a two-tailed unpaired student t-test was performed between two groups.

## Data and materials availability

All the data and relevant materials, including reagents and primers, that supports the findings of this study are available from the corresponding author upon reasonable request.

## Acknowledgements

We thank Yanling Yan (Huazhong University of Science and Technology, China) for critical discussions. We thank the Mass Spectrometry Core Facility (Mr. Cookson K. C. Chiu) and the Bio-imaging Core Facility (Dr. Zhenglong Sun and Ms. Mei Yu) of Shenzhen Bay Laboratory for providing technical supports.

## Author contributions

T. Zhou. and W. Ji. conceived the project and designed the experiments. T. Zhou. and Y. Du performed the experiments T. Zhou., Y. Du., X. Hu., A. Shi., W. Chang., L. Deng. And W. Ji. analyzed and interpreted the data. W. Ji. prepared the manuscript with inputs and approval from all authors.

---

**Supplementary Fig. 1.**
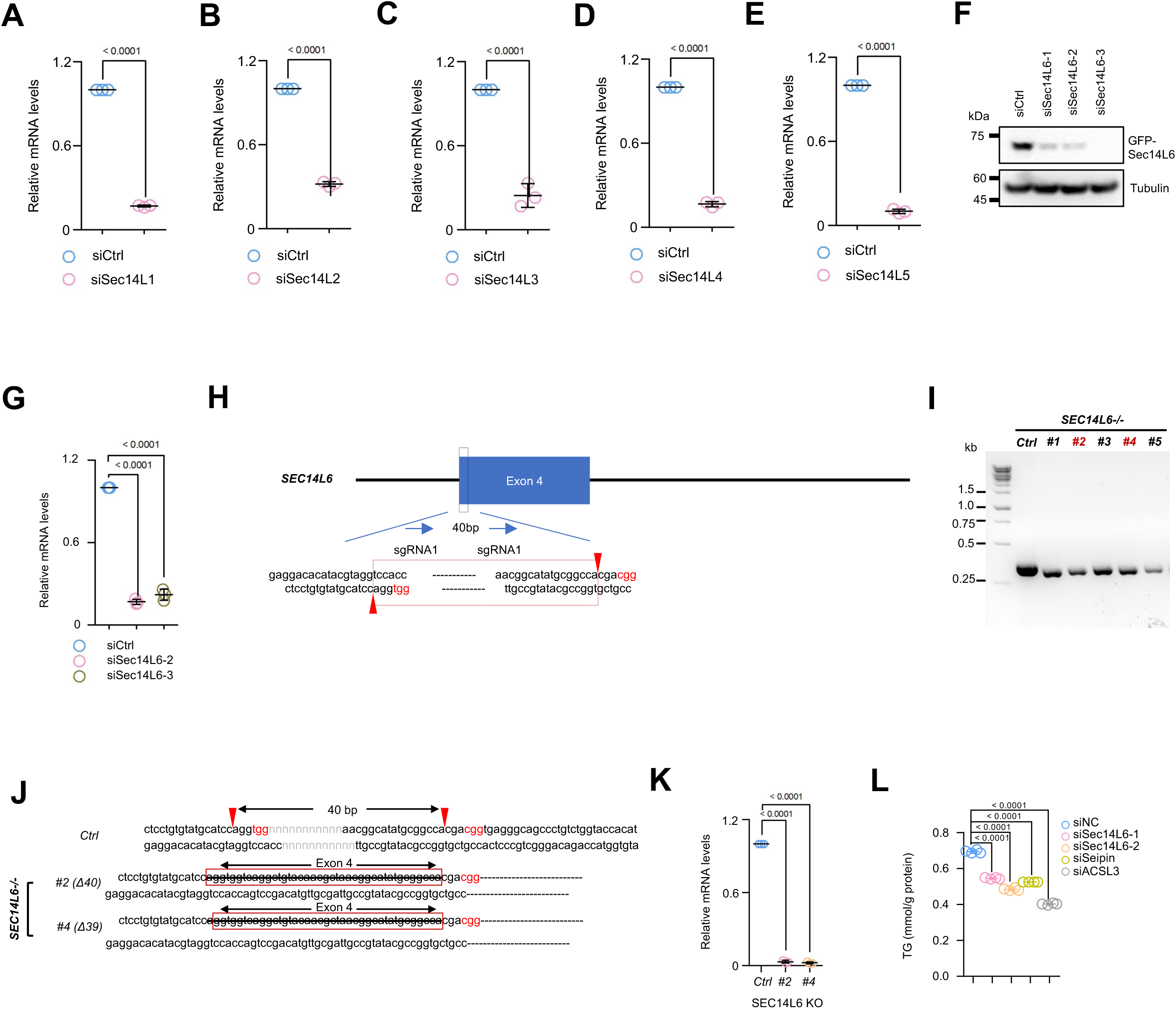
Supplemental results to Fig.1. **A-E**. qPCR showed the efficiency of siRNA-mediated suppression of Sec14L1 (**A**), Sec14L2 (**B**), Sec14L3 (**C**), Sec14L4 (**D**) or Sec14L5 (**E**) in more than 3 independent experiments. Two-tailed unpaired student t-test. Mean ± SD. **F.** Immunoblots showing the efficiency of Sec14L6 siRNAs used in Fig.1. **G.** qPCR showed the mRNA levels of Sec14L6 in siRNA-mediated Sec14L6 suppression as Fig.1 in 3 independent experiments. Ordinary one-way ANOVA with Tukey’s multiple comparisons test. Mean ± SD. **H.** Schematic diagram of CRISPR-CAS9-mediated KO of Sec14L6 in HeLa cells. **I, J.** Two Sec14L6 clones (#2 and #4) were confirmed by DNA gels (**I**) and sequencing (**J**). In (H), red rectangles highlight the deleted region of Sec14L6-KO #2 and Sec14L6-KO #4. **K**. qPCR showed the mRNA levels of Sec14L6 in Sec14L6-KO #2 and Sec14L6-KO #4 in 3 independent experiments. Ordinary one-way ANOVA with Tukey’s multiple comparisons test. Mean ± SD. **L**. Cellular levels of TG in HeLa cells treated with Sec14L6, Seipin or ACSL3 siRNAs from more than 3 independent experiments. Ordinary one-way ANOVA with Tukey’s multiple comparisons test. Mean ± SD.

**Supplementary Fig. 2.**
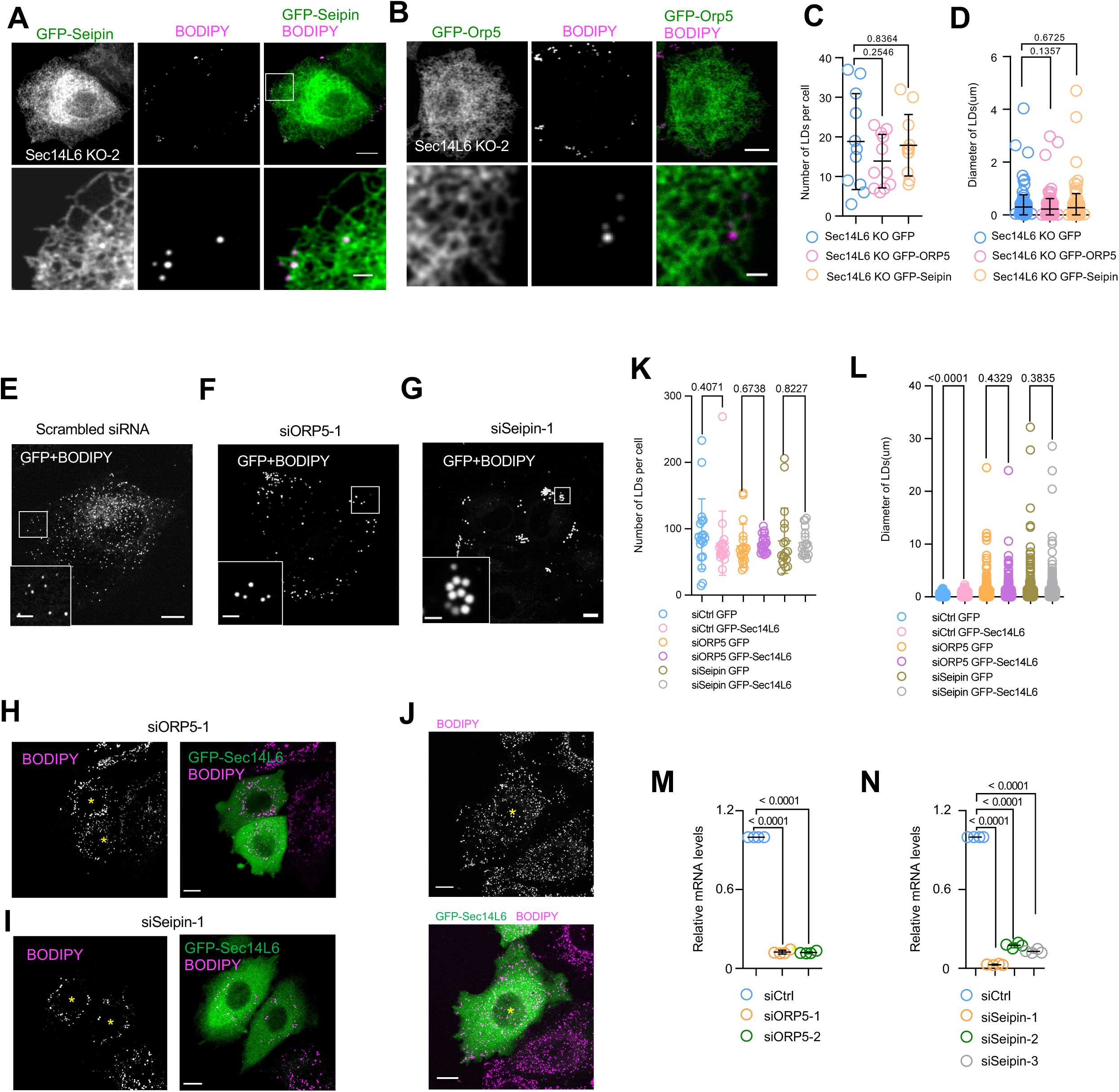
The functional relationship between Sec14L6 and known LD biogenesis factors Seipin and Orp5. **A, B**. Representative images of live BODIPY (magenta)-stained Sec14L6 KO HeLa cells (KO-2) transiently expressing GFP-Seipin (green, **A**) or GFP-Orp5 (green, **B**) with insets. **C, D**. The number (**C**) or size (**D**) of LDs in cells as shown in (**A, B**) in more than 3 independent experiments. Ordinary one-way ANOVA with Tukey’s multiple comparisons test. Mean ± SD. **E-I**. Representative images of live BODIPY (magenta)-stained HeLa cells transiently expressing GFP or GFP-Sec14L6 (green) upon control (**E**), Orp5 (**F, H**) or Seipin siRNAs (**G, I**) with insets. **J**. Representative images of live BODIPY (magenta)-stained HeLa cells transiently expressing GFP-Sec14L6 (green). Yellow asterisks denote a cell with GFP-Sec14L6 expression. **K, L.** The number (**K**) or size (**L**) of LDs in cells as shown in (**E-J**) in more than 3 independent experiments. Ordinary one-way ANOVA with Tukey’s multiple comparisons test. Mean ± SD. **M, N.** qPCR showing the efficiency of suppression of Orp5 (**F, H**) and Seipin (**G, I**) in cells as in (**E-I**) from 3 independent experiments. Ordinary one-way ANOVA with Tukey’s multiple comparisons test. Mean ± SD. Scale bar, 10μm in the whole cell images and 2μm in the insets in (A, B & E-I).

**Supplementary Fig. 3.**
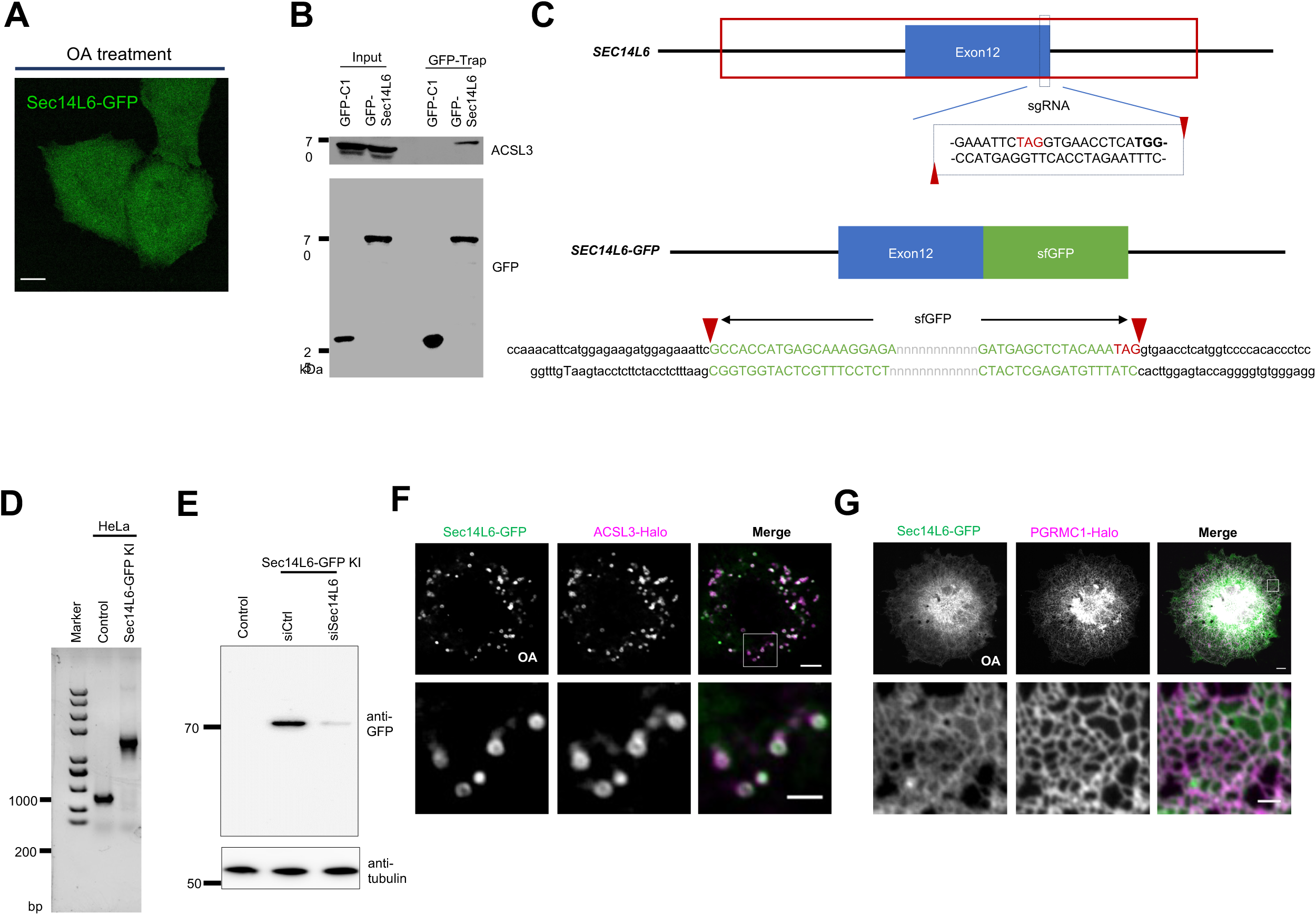
Supplemental results to Fig.2. **A.** Representative images of live HeLa cells transiently expressing Sec14L6-GFP (green). **B.** GFP-Trap assays demonstrated an interaction between GFP-Sec14L6 and endogenous ACSL3. **C.** Schematic diagram of CRISPR-CAS9-mediated Knockin of GFP at the C terminus of the Sec14L6 gene in HeLa cells. **D.** DNA gel of the Sec14L6-GFP-KI line. **E.** Immunoblots of the Sec14L6-GFP-KI line treated with control or Sec14L6 siRNAs using GFP antibody. **F, G**. Representative images of live HeLa cells transiently expressing Sec14L6-GFP (green) along with ACSL3-Halo (**F**, magenta) or PGRMC1-Halo (**G**, magenta) with insets on the bottom. Scale bar, 10μm in the whole cell images in (A, F & G).

**Supplementary Fig. 4.**
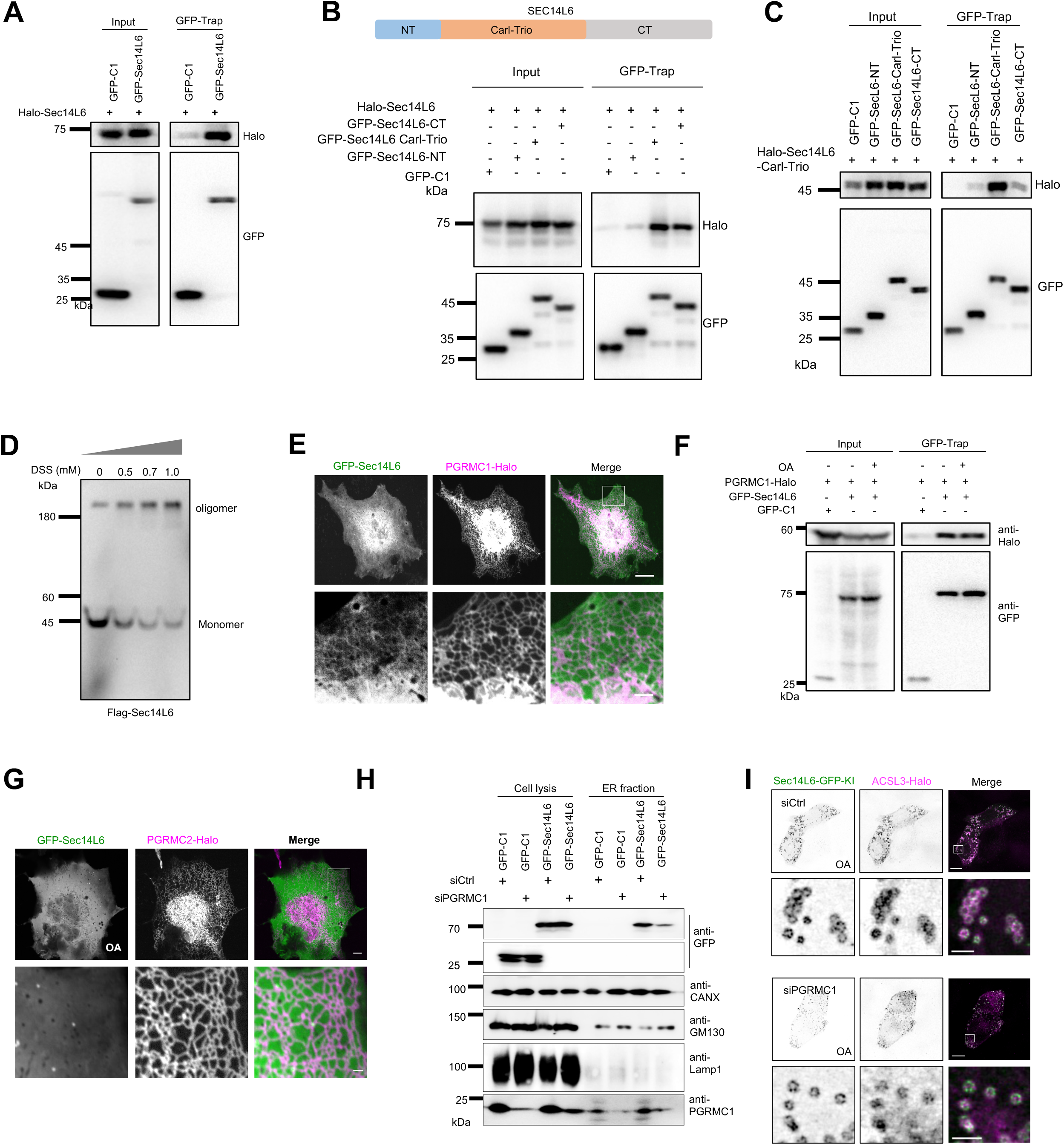
Supplemental results to Fig.3. **A.** GFP-Trap assays demonstrated self-interactions between GFP-Sec14L6 and Halo-Sec14L6. **B.** GFP-Trap assays demonstrated Sec14L6 self-interactions occurred between GFP-Sec14L6 and the Carl-Trio domain or the CT region. **C.** GFP-Trap assays demonstrated strong interactions among the Carl-Trio domain. **D.** Immunoblots of flag-Sec14L6 upon cross-linker DSS treatments (0, 0.5, 0.7, 1.0 mM) in HeLa cells. **E.** Representative images of live HeLa cells transiently expressing GFP-Sec14L6 (green) and PGRMC1-Halo (magenta) in absence of OA with an inset. **F.** GFP-Trap assays demonstrated interactions between GFP-Sec14L6 and PGRMC1-Halo in absence or prence of OA. **G.** Representative images of live HeLa cells transiently expressing GFP-Sec14L6 (green) and PGRMC2-Halo (magenta) under OA treatment with an inset. **H.** Immunoblots showed the level of GFP-Sec14L6 in ER membrane fractions of either control or PGRMC1-depleted cells stably using GFP vector as a negative control. **I.** Representative images of live Sec14L6-GFP-KI (green) HeLa cells transiently expressing ACSL3-Halo (magenta) upon scrambled or PGRMC1 siRNA treatments after OA loading (3 h) with insets on the bottom. Scale bar, 10μm in the whole cell image and 2μm in inset (E, G & I).

**Supplementary Fig.5.**
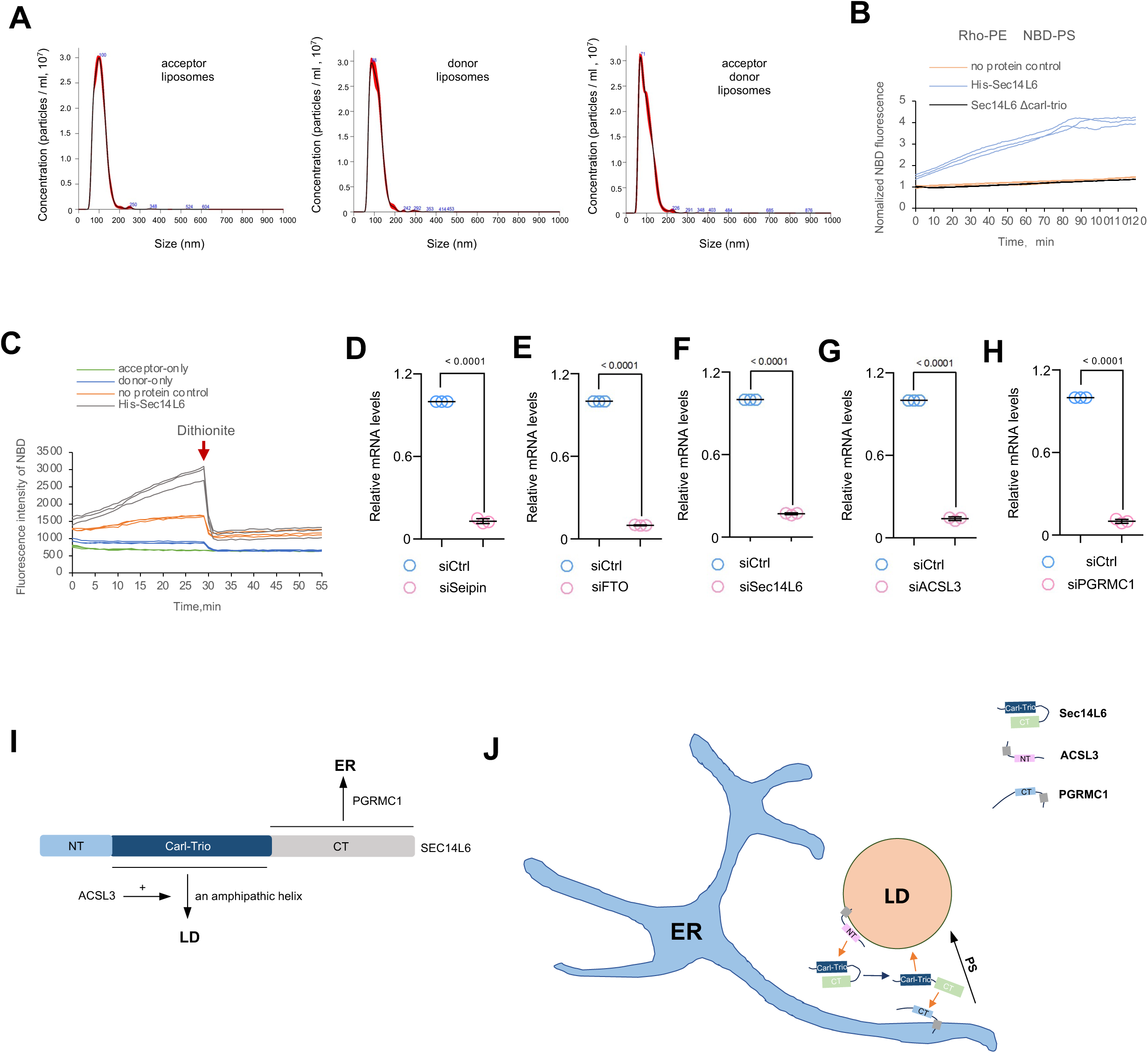
Supplementary results related to Fig. 6-9. **A.** Particle analyses showed the distribution of the size of donor and acceptor liposomes used in this study. **B.** As in Fig.6I, donor liposomes (20 μM) containing fluorescent lipids [2% NBD-PS, 2%, 2% Rhodamine-PE, 5% DGS-NTA, 61% DOPC, and 30% PE) were mixed 1:1 with acceptor liposomes [20 μM: 65% DOPC, 30% PE, and 5% DGS-NTA] in the absence or presence of His-Sec14L6 proteins (0.25 μM). The measurements lasted for 120 min and the NBD fluorescence was normalized to the fluorescence intensity at time=0. **C.** The dithionite assay demonstrated that His-Sec14L6 did not promote liposome fusion. In this assay, dithionite solution was added to the liposomes after the lipid transfer reactions as shown in Fig. 6H. The NBD fluorescence was normalized to the fluorescence intensity at time=0. **D-H.** qPCR showing the efficiency of suppression of Seipin (**D**), FTO (**E**), Sec14L6 (**F**), ACSL3 (**G**) and PGRMC1 (**H**) in cells as in Fig.9 from 3 independent experiments. Two-tailed unpaired student t-test. Mean ± SD. **I.** The membrane association of Sec14L6 and related mechanisms identified in this study. **J.** Working model of Sec14L6 in LD biogenesis. We propose a model, in which Sec14L6 transfers PS and/or PI4P between the ER and LDs to promote LD formation. Sec14L6 interacts with ACSL3 and the interaction facilitates the association of Sec14L6 with LDs, and stimulates the PS transfer activity of Sec14L6 in vitro. PGRMC1 acts as an adaptor that recruits of Sec14L6 to the ER.

**Video 1**. Representative time-lapse images of a HeLa cell transiently expressing GFP-Sec14L6 (green) and ACSL3-Halo (magenta) under EBSS-OA condition. Left: ACSL3-Halo; right: merged image. (time interval: 2 second; frame rate: 5 frame per second).

